# Viridiplantae-specific GLXI and GLXII isoforms co-evolved and detoxify glucosone *in planta*

**DOI:** 10.1101/2022.09.13.507747

**Authors:** Manuel Balparda, Jessica Schmitz, Martin Duemmel, Isabell C. Wuthenow, Marc Schmidt, Saleh Alseekh, Alisdair R. Fernie, Martin J. Lercher, Veronica G. Maurino

**Affiliations:** Molekulare Pflanzenphysiologie, Institut für Zelluläre und Molekulare Botanik, Rheinische Friedrich-Wilhelms-Universität Bonn, Kirschallee 1, 53115 Bonn, Germany; Plant Molecular Physiology and Biotechnology, Institute of Developmental and Molecular Biology of Plants, Heinrich Heine University, 40225 Düsseldorf, Germany; Max-Planck-Institute of Molecular Plant Physiology, Am Mühlenberg 1, 14476 Potsdam-Golm, Germany; Center for Plant Systems Biology and Biotechnology, 4000 Plovdiv, Bulgaria; Institute for Computer Science and Department of Biology, Heinrich Heine University, 40225 Düsseldorf, Germany

**Author notes:** Both authors contributed equally to the work (Manuel Balparda, Jessica Schmitz). Corresponding author: Veronica G. Maurino. The author responsible for distribution of materials integral to the findings presented in this article is: Veronica G. Maurino.

## Abstract

Reactive carbonyl species (RCS) such as methylglyoxal (MGO) and glyoxal (GO) are highly reactive, unwanted side-products of cellular metabolism maintained at harmless intracellular levels by specific scavenging mechanisms. MGO and GO are metabolized through the glyoxalase (GLX) system, which consists of two enzymes acting in sequence, GLXI and GLXII. While plant genomes encode a large number of different GLX isoforms, it is unclear what their specific functions are and how these arose in evolution. Here, we show that plants possess two GLX systems of different evolutionary origins and with distinct structural and functional properties. The first system is shared by all eukaryotes, scavenges MGO and GO especially during seedling establishment, and features Zn^2+^-type GLXI, a metal co-factor preference that arose already in the last eukaryotic common ancestor. The GLXI and GLXII of the second system can together metabolize KDG, a glucose-derived RCS, and were acquired by the last common ancestor of viridiplantae through horizontal gene transfer from proteobacteria. In contrast to bacterial GLXI homologs, which are active as dimers, plant Ni^2+^-type GLXI contain a domain duplication, are active as monomers, and have modified their second active site. The acquisition and neofunctionalization of a structurally, biochemically, and functionally distinct GLX systems indicate that viridiplantae are under strong selection to detoxify a diversity of RCS.

## Introduction

The ecological success of plants is based on the evolution of mechanisms they use to adapt to environmental changes. Under stress, and to a lesser degree also during normal conditions of growth, plant metabolism produces reactive oxygen species (ROS) and reactive carbonyl species (RCS), small molecules that are toxic due to their high reactivity towards cellular macromolecules. To limit their deleterious effects, plants evolved scavenging systems, which convert ROS and RCS into less harmful molecules that feed into intermediary metabolism (Hanson et al., 2016; Hudig et al., 2018). RCS are small electrophilic mono- and di-carbonyl molecules, which are highly reactive toward cellular macromolecules. They are potent glycating agents of proteins, nucleotides and basic phospholipids, directly contributing to cell damage through these glycation processes (Murata-Kamiya et al., 2000; Thornalley, 2003; Sibbersen et al., 2013; Sousa Silva et al., 2013; Vistoli et al., 2013; Semchyshyn, 2014) or via the formation of advanced glycation products (AGEs) (Thornalley, 2008). In humans, elevated levels of RCS and AGEs are involved in aging and in several degenerative processes and diseases, including diabetes as well as neurodegenerative and cardiovascular diseases (Goldin et al., 2006; Thornalley, 2008; Munch et al., 2012). In plants, RCS accumulate under various abiotic stresses such as salinity, drought, and cold (Yadav et al., 2005). Furthermore, upon exposure to high light or elevated CO_2_ concentration, AGEs formed by enhanced RCS levels were found to accumulate in leaves (Qiu et al., 2008; Bechtold et al., 2009).

The most abundant RCS in plant tissues are the alpha-oxoaldehydes (1,2 di-carbonyls) glyoxal (GO) and methylglyoxal (MGO). Sources of GO and MGO in plant metabolism are lipid peroxidation, the fragmentation of glycated proteins, and the auto-oxidation of glucose (Thornalley et al., 1999; Yin and Porter, 2005). The main pathway of MGO formation is glycolysis, where the enediolate phosphate reaction intermediate of the triose phosphate isomerase reaction can be non-enzymatically decomposed (Richard, 1991; Phillips and Thornalley, 1993). In addition, the participation of triose phosphate isomerase in the Calvin-Benson cycle adds a source of MGO during photosynthesis (Chen and Thelen, 2010; Takagi et al., 2014). Two other highly reactive alpha-oxoaldehydes are keto-d-glucose (glucosone; KDG) and 3-deoxyglucosone (3-DG), which arise as byproducts of sugar metabolism (Bean and Hassid, 1956; Baute, 1984; Semchyshyn, 2014) through the oxidation of d-glucose (Thornalley et al., 1999) and the cleavage of glycated proteins (Nagai et al., 2002).

The main pathway to scavenge GO and MGO is the glyoxalase (GLX) two-enzyme system. The first enzyme, *S*-d-lactoylglutathione lyase (glyoxalase I, GLXI), converts hemithioacetals formed by the non-enzymatic reaction of alpha-oxoaldehydes with glutathione (GSH) (e.g., MGO-GSH or GO-GSH) to an *S*-2-hydroxyacylglutathione intermediate. The second enzyme, *S*-2-hydroxyglutathione hydrolase (glyoxalase II, GLXII), converts the product of the GLXI reaction into a (D)-α-hydroxyacid derivative (Thornalley, 1990; Schmitz et al., 2017), glycolate in the case of GO and d-lactate in the case of MGO (Schmitz et al., 2017). Glycolate can be further metabolized by peroxisomal glycolate oxidase (Maurino and Engqvist, 2015), and d-lactate is converted to pyruvate through mitochondrial d-lactate dehydrogenase (Engqvist et al., 2009; Welchen et al., 2016). While the GLX system’s role for the detoxification of GO and MGO is well established, it is unclear whether the same system can protect plant cells from KDG or 3-DG toxicity.

In *Arabidopsis thaliana*, the GLX system is genomically encoded by three homologs of *GLXI* (GLXI;1, GLXI;2 and GLXI;3) and three homologs of *GLXII* (GLXII;2, GLXI;4 and GLXI5) (Maiti et al., 1997; Marasinghe et al., 2005; Schmitz et al., 2017), from which several GLXI and GLXII isoforms with different subcellular localization derive through alternative splicing (Schmitz et al., 2017).

GLXI is a metalloenzyme, and is customarily classified into Ni^2+^- and Zn^2+^-type proteins (Sukdeo et al., 2004). In phylogenies, GLXI;1 and GLXI;2 cluster with known Ni^2+^-dependent GLXI, such as *E. coli* GLXI (BAE76494); we thus classify them as Ni^2+^-type GLXI. In contrast, GLXI;3 clusters with known Zn^2+^-dependent GLXI such as human GLXI (AAD38008) (Schmitz et al., 2017; Schmitz et al., 2018), and we hence classify it as Zn^2+^-type GLXI. Upon close inspection, Arabidopsis GLXI isoforms defy such a simple classification, however, as they can use other metal ion cofactors and can change their substrate specificities depending on the metal cofactor used. While GLXI;3 strongly prefers Mn^2+^ in the conversion of both MGO-GSH and GO-GSH, GLXI;1 and GLXI;2 prefer Ni^2+^ in the conversion of MGO-GSH and Mn^2+^ in the conversion of GO-GSH (Schmitz et al., 2017). The Arabidopsis GLXI isoforms have similar affinities towards MGO-GSH and GO-GSH, but exhibit different kinetic behaviors. GLXI;3 has a much higher catalytic efficiency for MGO-GSH and GO-GSH than GLXI;1 and GLXI;2, and is the isoform mostly involved in detoxification of MGO during germination and post-germinative development (Schmitz et al., 2017). The second enzyme of the pathway, GLXII, belongs to the beta-lactamase protein family and contains a Fe^3+^/Zn^2+^ binuclear metal center (Cricco and Vila, 1999; Crowder and Walsh, 1999; Zang et al., 2001; Wenzel et al., 2004; Marasinghe et al., 2005). GLXII are active as dimers, which can act on a broad spectrum of glutathione thioesters (Thornalley, 1990; Ridderstrom and Mannervik, 1996; Bito et al., 1997; Marasinghe et al., 2005).

What could be the evolutionary advantages of harboring multiple GLXI and GLXII isoforms? Gene duplication followed by sub-functionalization or neo-functionalization is a potent driving force in the evolution of biological diversity, allowing functional divergence and innovation (Panchy et al., 2016; Kuzmin et al., 2022). It is believed that multiple isoforms of a plant enzyme typically act on alternative substrates, or on the same substrate with distinct kinetics optimized for different conditions or cellular compartments (Sappl et al., 2004; Maurino et al., 2009). It thus appears likely that the existence of GLXI and GLXII isoforms with different localization and biochemical properties in plant cells resulted from the evolution of enzymatic variants optimized for the metabolization of alternative RCS produced in specific plant metabolic pathways, subcellular compartments, or growth conditions.

Unfortunately, very little is known about the evolution of the plant GLX system. The phylogenetic relationships of plant GLXII proteins have not been studied at all. For GLXI proteins, only one report exists (Kaur et al., 2013), which claims to elucidate episodes of horizontal gene transfer and gene fusion that led to GLXI diversification. However, the main conclusions of this study are not supported by the evidence it presents (see Suppl. Text), and hence the evolution of the GLX system in plants is still an open question.

At the molecular functional level, the plant GLX system was only studied as a scavenging system for MGO or GO (Thornalley, 1990; Schmitz et al., 2017), although it appears likely that it also plays a role in the detoxification of other alpha-oxoaldehydes produced by cellular metabolic activities. A full understanding of the functions and biochemical properties of the GLX system is also of practical importance, as this scavenging system is included in current strategies that aim to improve plant fitness and yield by mitigating abiotic stresses (Alvarez Viveros et al., 2013; Sankaranarayanan et al., 2017).

Here, we aim to clarify the evolutionary history of GLXI and GLXII in plants and to shed light on the physiological advantages for plants to retain multiple GLXI and GLXII isoforms. For this, we first analyzed the evolutionary emergence of GLXI and GLXII proteins, building phylogenies for comprehensive sequence sets that span the tree of life. These analyses confirm the existence of GLXI and GLXII variants that co-existed in the last eukaryotic ancestor and that persist in all major eukaryotic lineages. In addition, we found that viridiplantae possess a unique two-domain GLXI, which likely arose through the duplication of a GLXI sequence likely acquired together with the ancestral sequence of viridiplantae-specific GLXII via horizontal gene transfer from a proteobacterium. Amino acid changes in the second structural domain of viridiplantae-specific GLXI led to the formation of two unequal binding sites, suggesting a functional specialization of these isoforms. Using *Arabidopsis thaliana* as model species and a combination of biochemical and physiological approaches, we demonstrate that the viridiplantae-specific GLXI and GLXII isoforms are involved in the metabolization of KDG in vivo. It appears that early on in its evolution, the green lineage developed a unique solution to detoxify KDG, for which it evolved a parallel GLX system, derived from a pair of gene sequences acquired from a proteobacterium.

## Results

### Two-domain GLXI and a monophyletic cluster containing organellar-localized GLXII are unique to viridiplantae

We assembled sets of potential GLXI and GLXII sequences from representative species across the tree of life by blasting protein sequences from five representative species (*Homo sapiens, Arabidopsis thaliana, Phaeodactylum tricornutum, Synechocystis salina*, and *Escherichia coli*) against all sequences in RefSeq (Pruitt et al., 2012) (see Methods; results in Supplemental File 1 and 2). Some GLXI homologs form dimers, where each monomer builds one structural domain, while other GLXI homologs are monomers containing two similar structural domains (Ridderstrom and Mannervik, 1996; Frickel et al., 2001). We split all two-domain sequences into structural domains A and B at the hinge region (Supplemental Fig. 1). We then built maximum-likelihood gene trees for the GLXI and GLXII sequence sets.

Our phylogenetic analysis indicates that plant homologs of GLXI proteins can be grouped into two clades of separate evolutionary origin (Fig. 1A and Supplemental Fig. 2). The GLXI sequences, commonly referred to as Zn^2+^-type GLXI and represented by GLXI;3 in Arabidopsis, are conserved across eukaryotic kingdoms. The topology of the gene tree (Fig. 1a and Supplemental Fig. 2) indicates that a copy of this protein was present in the last eukaryotic common ancestor (Fig. 1B). The closest prokaryotic relatives of the eukaryotic Zn^2+^-type GLXI are found in proteobacteria (Fig. 1a and Supplemental Fig. 2), consistent with the likely origin of eukaryotes through the endosymbiosis of an ancestral α-proteobacterium in an archaeon. In contrast, the GLXI sequences commonly referred to as the Ni^2+^-type class – represented by GLXI;1 and GLXI;2 in Arabidopsis, and distinguished from the Zn^2+^-type form through conserved amino acids (see next subsection) – are only present in proteobacteria, cyanobacteria, and viridiplantae (streptophyta + chlorophyta) (Fig. 1A and Supplemental Fig. 2). The topology of the gene tree indicates that the Ni^2+^-type GLXI was likely acquired by the common ancestor of viridiplantae through horizontal gene transfer from a proteobacterium (Fig. 1b). Thus, viridiplantae are the only eukaryotes that possess two classes of GLXI with different evolutionary origins. Arabidopsis GLXI;1 possesses a long amino terminal sequence (Supplemental Fig. 3), which was shown to direct the protein to chloroplasts (Schmitz et al., 2017). Interestingly, all homologs of viridiplantae GLXI;1 also possess a long amino terminal sequence (Supplemental Fig. 3), indicating that these proteins are most likely also localized to organelles; the only exception is one protein in rice, which lacks a long terminal sequence.

**Figure 1.**
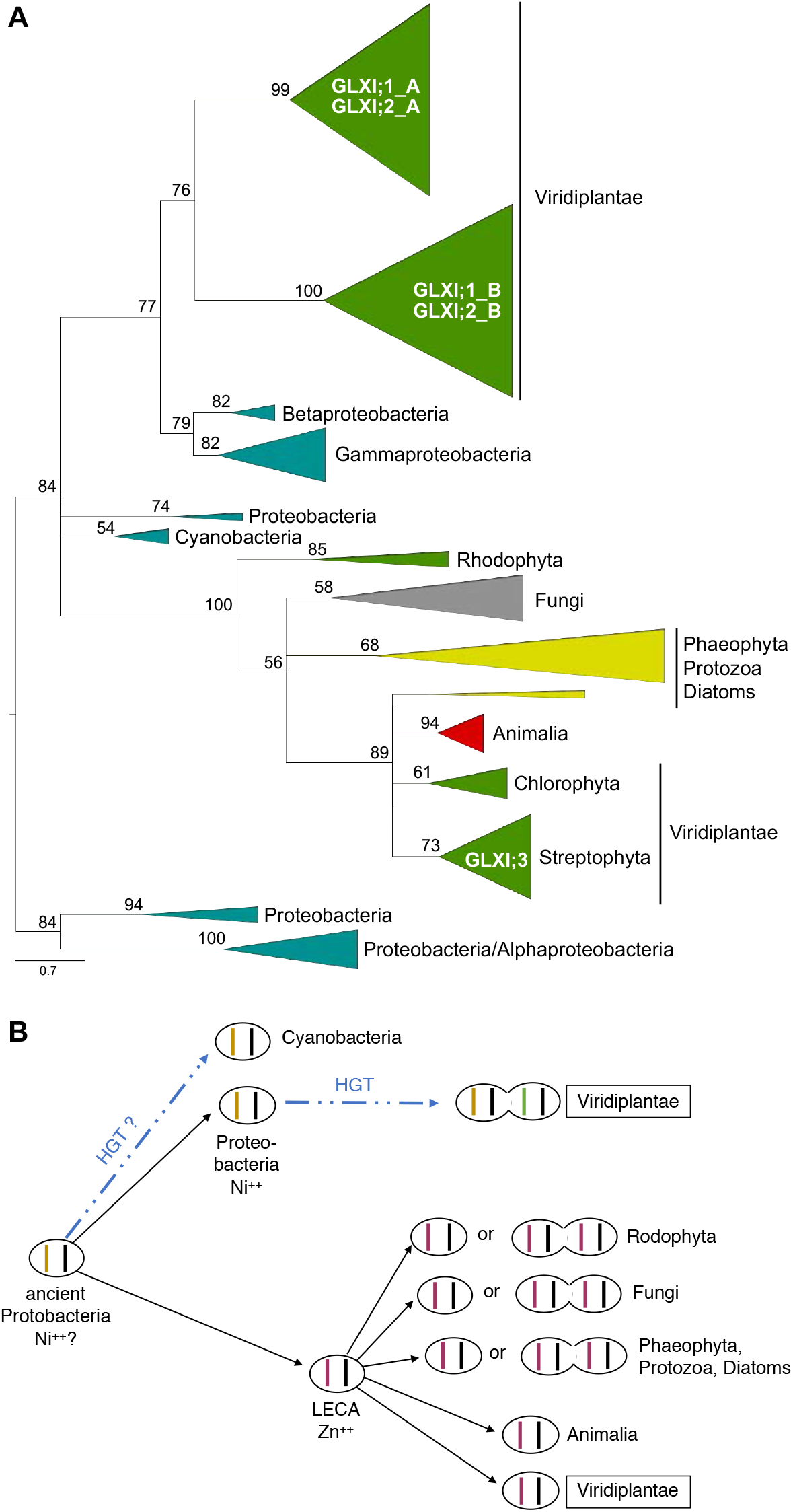
Evolutionary analysis puts plant GLXI into two clades grouping with other eukaryotes and proteobacteria, respectively. **A**, Summary of the evolutionary history of GLXI domains, inferred using Maximum Likelihood. We split all two-domain GLXI sequences at the position corresponding to E146 in *A. thaliana* GLXI;2 into A and B parts before aligning them. Numbers atop branches show percent bootstrap support values from 1000 replicates. For this summary tree, branches with BS<50 were collapsed into polytomies, and triangles group related organisms. Branch lengths reflect the number of substitutions per site. The full, uncollapsed tree with all individual sequences is shown in Supplemental Fig. 2. **B**, Depiction of the hypothesized evolutionary trajectories of glyoxalase systems. Single ovals represent systems with single-domain GLXI, double ovals represent systems with two-domain GLXI; the vertical bars inside indicate the putative GLXI active sites, with different colors indicating different amino acids at the conserved positions (see text).

Closely mirroring the situation for GLXI, homologs of GLXII also cluster in two clades of separate origin (Fig. 2 and Supplemental Fig. 4). Like Zn^2+^-type GLXI proteins, the class of GLXII represented by GLXII;2 in Arabidopsis is conserved across eukaryotic kingdoms (Fig. 2 and Supplemental Fig. 4), and the pattern of inheritance among eukaryotes is largely consistent with vertical transmission. Again, this indicates that this class represents the ancestral eukaryotic enzyme form, and that a copy of this protein was present in the last common ancestor of all eukaryotes. The closest bacterial relatives of this class are delta-proteobacteria, while the bacterial endosymbiont at the origin of eukaryotes is believed to have been more closely related to extant alpha-proteobacteria. However, it is still possible that this endosymbiont contributed the ancestral GLXII sequence, which it could have acquired previously through horizontal gene transfer from delta-proteobacteria.

**Figure 2.**
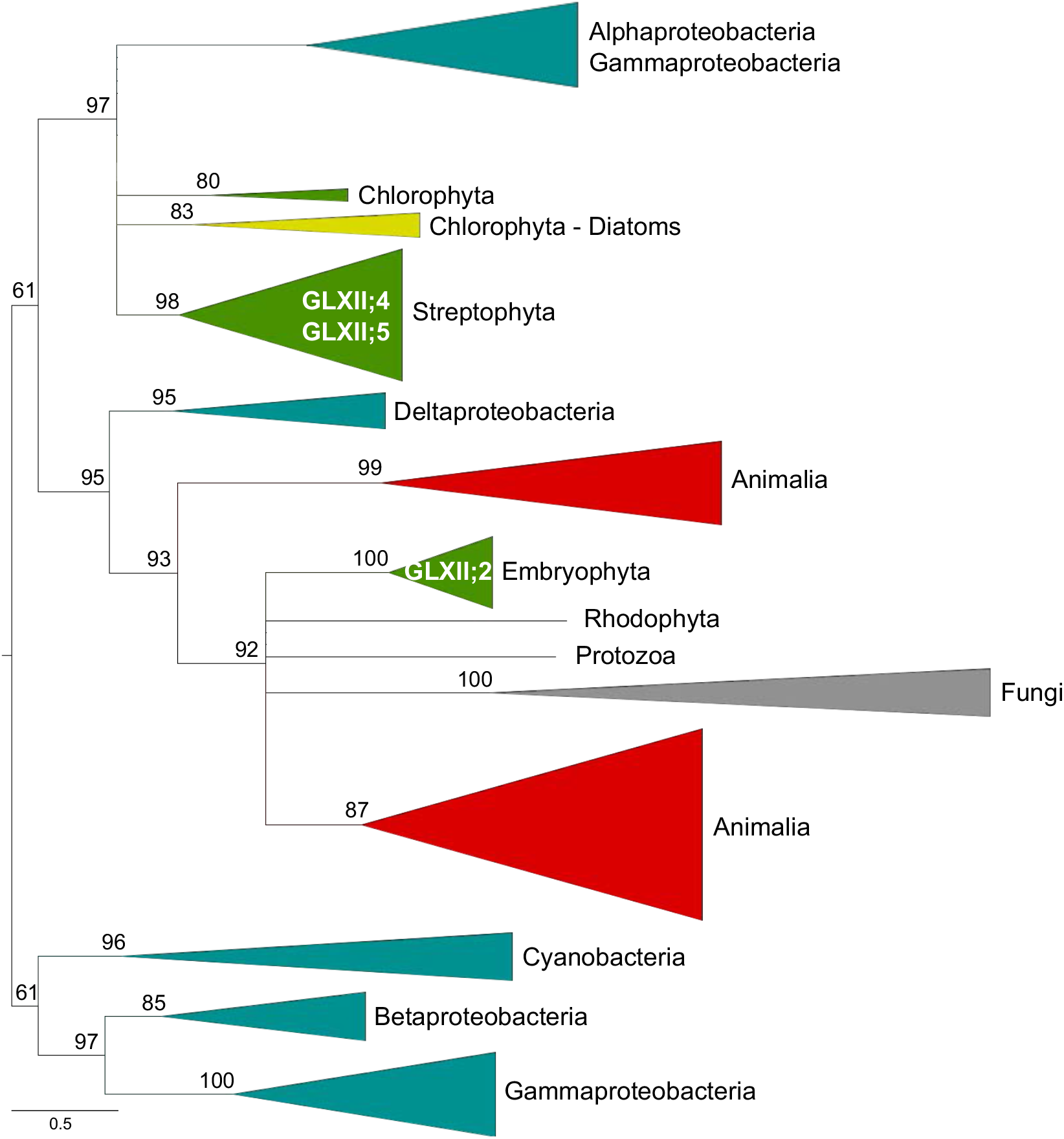
Evolutionary analysis puts plant GLXII into two clades grouping with other eukaryotes and proteobacteria, respectively. The figure summarizes the evolutionary history of GLXII sequences, inferred using Maximum Likelihood. Numbers atop branches show percent bootstrap support values from 1000 replicates. For this summary tree, branches with BS<50 were collapsed into polytomies, and triangles group related organisms. Branch lengths reflect the number of substitutions per site. The full, uncollapsed tree with all individual sequences is shown in Supplemental Fig. 3.

Similar to Ni^2+^-type GLXI proteins, the GLXII class proteins represented by GLXII;4 and GLXII;5 in Arabidopsis is found in proteobacteria and viridiplantae. The only difference is the presence of this class of proteins also in diatoms. However, this is not surprising, since diatoms arose through endosymbiosis involving a green alga (Moustafa et al., 2009; Deschamps and Moreira, 2012). Interestingly, many sequences of viridiplantae from this cluster possess a long terminal sequence (Supplemental Fig. 5), which was shown to direct the proteins to both chloroplasts and mitochondria in Arabidopsis GLXII;4 and GLXII;5 (Schmitz et al., 2017). This observation suggests that in the green lineage, this class of GLXII tends to be active in organelles. As in the case of the Ni^2+^-type GLXI proteins, the most likely scenario for the acquisition of this class of GLXII by viridiplantae is gene transfer from a proteobacterium to the common ancestors of the green lineage, possibly involving a co-transfer of GLXII and the ancestor of Ni^2+^-type GLXI genes.

### Amino acid conservation analysis reveals the evolution of specific GLXI isoforms in viridiplantae

Dimers of single-domain GLXI as well as monomers of two-domain GLXI display the typical fold defining the GLXI family (Supplemental Fig. 1) (Kelley et al., 2015; Turra et al., 2015). In the dimeric enzymes, two equivalent active sites (active site 1 and 2, Fig. 3A) are formed by regions contributed by both monomers, which are arranged in an antiparallel fashion. Monomeric enzymes from two-domain homologs fold in a similar way, forming two active sites with contributions of both structural domains (active site 1 and 2, Fig. 3B). Our results show that Zn^2+^-type GLXI proteins are one-domain proteins in animalia and viridiplantae, whereas other groups, such as rhodophyta, phaeophyta, diatoms, protozoa, and fungi contain one as well as two-domain Zn^2+^-type GLXI proteins (Fig. 1B). Within each of these groups, the one- and two-domain sequences together form a monophyletic cluster (Supplemental Fig. 2), indicating independent domain duplications in different eukaryotic lineages. We found that in bacteria, Ni^2+^-type GLXI is a one-domain protein, while in all viridiplantae it is a two-domain protein containing the A and B structural domains (Fig. 1A and 3). This observation indicates that a domain duplication within the Ni^2+^-type GLXI gene occurred already in the last common ancestor of the green lineage.

**Figure 3.**
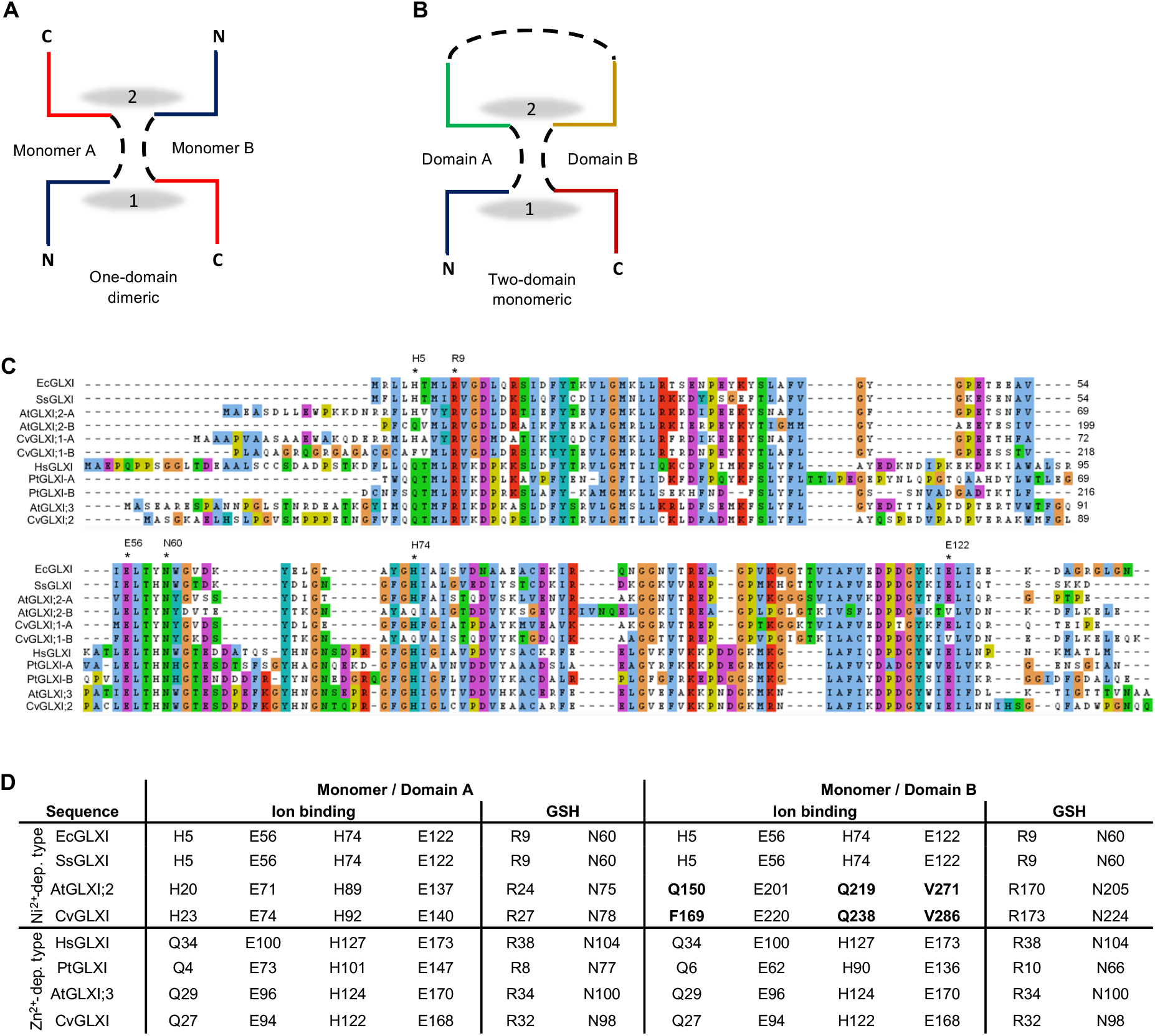
Analysis of GLXI proteins. **A**, Schematic representation of the folding of a canonical one-domain GLXI with formation of a homodimer. **B**, Schematic representation of the folding of a canonical monomeric two-domain GLXI. In A and B the folding of the proteins allow the formation of two similar active sites (1 and 2), with participation of amino acids of both monomers/domains. **C**, Alignment of representative GLXI proteins that group with known Ni^2+^-and Zn^2+^-dependent GLXI type proteins. The sequences of two-domain GLXI (AtGLXI;2, CvGLXI, and PtGLXI) were split at the amino acid corresponding to E146 in *A. thaliana* GLXI;2 into domain A and domain B. **D**, Conserved amino acid positions for the ion and GSH binding in GLXI proteins. Ec: *Echerichia coli*; Ss: *Synechocistis sp*; At: *A thaliana*; Cv: *Chlorella variabilis*; Hs: *Homo sapiens*; Pt: *Phaeodactylum tricornutum*.

Both structural domains in dimeric (one-domain) and monomeric (two-domain) GLXI proteins contribute to the formation of the active sites (Turra et al., 2015). In the case of the dimeric GLXI, the active sites are each formed by amino acid residues belonging to the two monomers (labeled ‘A’ and ‘B’ in Fig. 3A), while in the case of monomeric GLXI, the active sites are formed by amino acid residues belonging to corresponding structural domains A and B (Fig. 3B). We analysed the amino acid positions known to participate in the metal-ion and GSH binding in the active sites 1 and 2 of representative GLXI proteins (Fig. 3C) (He et al., 2000; Suttisansanee and Honek, 2011; Suttisansanee et al., 2015). We found that all Zn^2+^-type GLXI proteins possess the same amino acids at the indicated positions, independent of whether the active enzymes are dimers of one-domain proteins (represented by *Homo sapiens, A. thaliana* and *Chlorella variabilis* in Fig. 3C and D) or monomers of two-domain proteins (represented by domains A and B of *Phaeodactylum tricornutum* in Fig. 3C and D). This observation indicates that both active sites are functionally equivalent, as for the metal-ion binding in active site 1, the amino acids Q and E belong to monomer/structural domain A, and H and E belong to monomer/structural domain B (Fig. 3A and D); this is inverted for active site 2. The conserved amino acids for the binding of GSH, R and N, also remain unchanged (Fig. 3D).

We encountered a different situation with the Ni^2+^-type GLXI proteins. One-domain bacterial sequences (represented by *E. coli* in Fig. 3c and d) have two equivalent binding sites, with the amino acids H, E, H, and E for the metal-ion binding, and R and N for GSH binding (Fig. 3C and d). Thus, only the first amino acid for metal-ion binding is different from the conserved Zn^2+^-type GLXI sequences. In viridiplantae (represented by *A. thaliana* and *C. variabilis* in Fig. 3C and D), while domain A of the Ni^2+^-type GLXI presents the same specific amino acids found in the one-domain bacterial sequences (Fig. 3C and D), domain B presents three amino acid substitutions at the ion-binding site (Fig. 3C and bold face in Fig. 3D). These amino acid changes in the second structural domain result in the formation of two unequal binding sites.

Whereas the active site 1 possesses H, E, Q, and V at the metal-ion binding region, the active site 2 possesses Q, E, H, and E. The conserved amino acids of the GSH binding sites, R and N, remain unchanged. In plantae, the substitutions found at the metal-ion binding site are highly conserved; in chlorophytes, while the first substitution is H to Q in *Coccomyxa subellipsoidea* and H to F in *Chlorella variabilis*, the other two substitutions are the same as in plantae (Fig. 3C and D). All the amino acid substitutions that occurred in domain B of viridiplantae drastically change the physicochemical properties of the amino acids. All these observations suggest that after acquisition of the Ni^2+^-type GLXI by the ancestor of viridiplantae, the combination of a domain duplication with the diversification of the two binding sites, must have contributed new biochemical features, such as the ability to metabolize alternative substrates or to use other catalytic mechanisms.

### 2-Keto-d-glucose affects development of GLXI;2 loss-of-function seedlings

The above analyses indicate that eukaryotic Ni^2+^-type GLXI proteins evolved exclusively in viridiplantae (Fig. 1), and they show molecular adaptations that have changed the structure of one of the active sites (Fig. 3D). Recently, we showed that the activity of the Arabidopsis Zn^2+-^ type enzyme, GLXI;3, is crucial for the detoxification of GO and MGO during germination and seedling establishment, while the Ni^2+^-type enzymes, GLXI;1 and GLXI;2, are not essential for the detoxification of these molecules at that developmental stage (Schmitz et al., 2017). We thus hypothesize that in plants, GLXI;1 and GLXI;2 may act *in vivo* on RCS other than GO and MGO.

Keto-D-glucose (glucosone; KDG) and 3-deoxyglucosone (3-DG) are two alpha-oxoaldehydes produced as byproducts of glucose metabolism and have chemical properties that make them putative substrates of GLXI (Bean and Hassid, 1956; Baute, 1984; Semchyshyn, 2014). It is known that exposure of cells to alpha-oxoaldehydes by addition of the compound to the growth media induces growth arrest and toxicity (Shinoda, 1994; Kang et al., 1996; Schmitz et al., 2017). Thus, to analyse if any of the Arabidopsis GLXI isoforms could be involved in the detoxification of 3-DG and/or KDG, we first performed toxicity analyses using our previously produced loss-of-function mutants for the GLXI isoforms (Schmitz et al., 2017). Apart of analyzing the single mutants, to fully eliminate the Ni^2+^-type proteins we also included in the analysis the *glxI;1-2 glxI;2* double mutant produced by genetic crosses of the individual mutants (Schmitz et al., 2017).

We found that 3-DG did not cause any detectable effects in the development of the plant lines analysed. In contrast, KDG specifically impairs seedling establishment of the *glxI;2* single mutant; in the presence of KDG, *glxI;2* presented poor development of the root system (Fig. 4). In the same conditions, similar impairment in seedling development was observed in *glxI;1-2 glxI;2*, while KDG did not affected the establishment of *glxI;1* and *glxI;3* seedlings (Fig. 4). As a control, we included germination in the presence of MGO (Schmitz et al., 2017), where MGO largely affects the germination and seedling establishment of *glxI;3*, while the other mutants behave like the wild type (Fig. 4).

**Figure 4.**
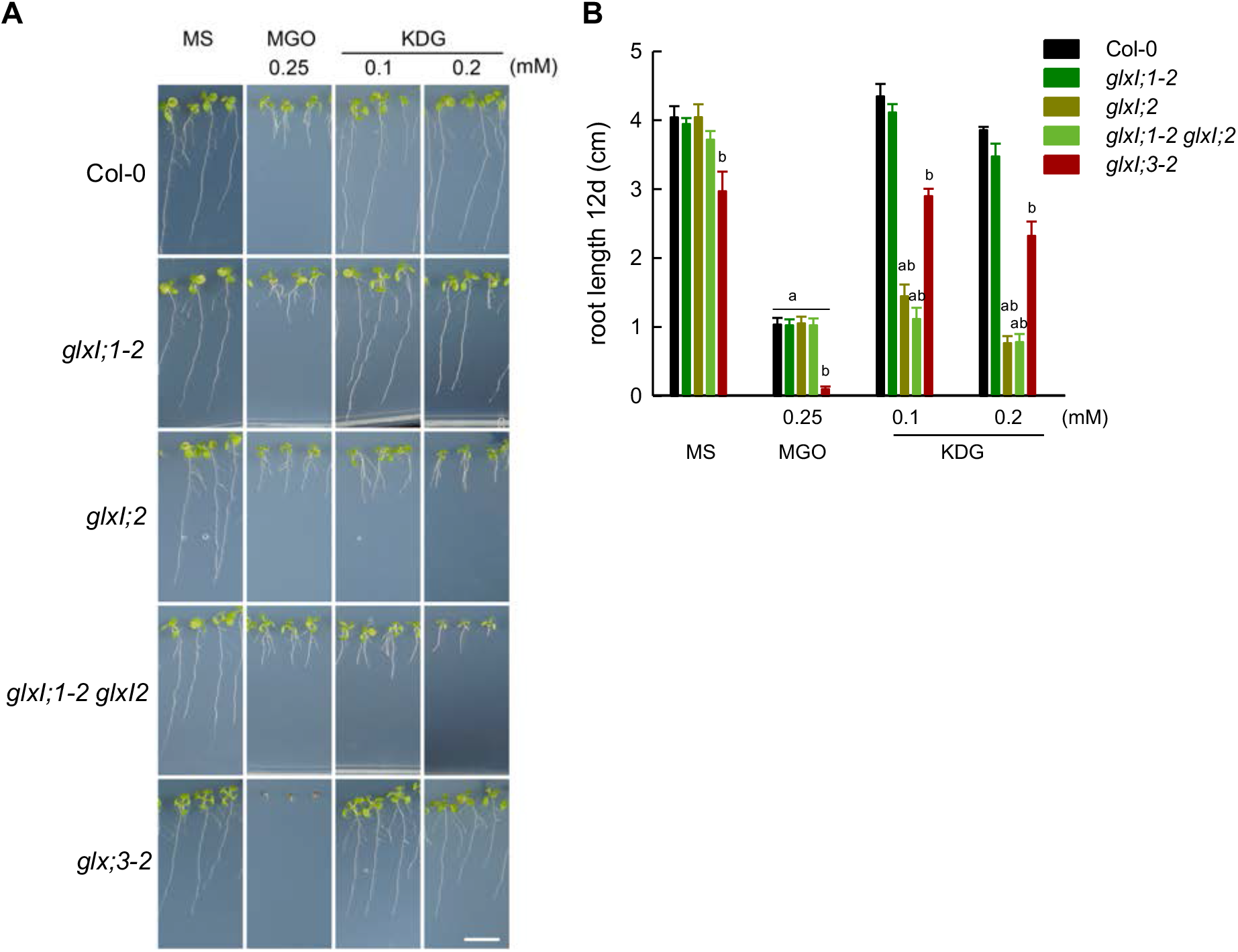
Effect of KDG and MGO on the seedling establishment of Arabidopsis GLXI T-DNA insertion lines. **A**, Representative picture of seedling development after 12 days of germination in MS medium and in the presence of MGO or KDG. Bar=1cm. **B**, Root length after 12 days of germination in MS medium and in the presence of MGO or KDG (as in A). Values are means ± SE from independent experimental trials (n=3-4) of 7-10 root measurements each. Testing for significant differences was performed by two-way ANOVA (treatment: P<0.0001, genotype: P<0.0001, interaction: P<0.0001). “a” and “b” indicate significant results of Tukey’s multiple comparison post-hoc test (P<0.001), where “a” indicates significant differences between treatment and MS control within a given genotype, and “b” indicates significant differences between genotype and Col-0 in a given treatment.

### GLXI;2 is involved in KDG metabolization after germination

To confirm that GLXI;2 is involved in KDG metabolism, we generated additional, independent loss-of-function lines for the GLXI;2 gene locus (*CC@glxI;2*) using the CRISPR/Cas9 system. We selected the line *CC@glxI;2-1* for further analysis, which presented a homozygous deletion in the GLXI;2 gene between exons 2 and 4 (Supplemental Fig. 6A-E). *CC@glxI;2-1* plants show the presence of low amounts of a shortened *GLXI;2* transcript (Supplemental Fig. 7A). In addition, we complemented the *glxI;2* T-DNA insertion line by expressing the full-length genomic sequence of GLXI;2 under the control of the constitutive AtUBQ10 promoter. We obtained different complemented lines (Supplemental Fig. 7B) and selected two lines, *glxI;2-GLXI;2-C4 (C4*) and *glxI;2-GLXI;2-C15 (C15*) for further analyses.

We analysed germination and post-germinative development of the wild type, both loss-of-function mutants *glxI;2* and *CC@glxI;2-1* (from here on referred to as *CC@glxI;2*), and the complemented lines C4 and C5 on MS medium supplemented with either MGO, GO, or KDG. Similar to *glxI;2*, the loss-of-function line *CC@glxI;2* showed an impairment of seedling establishment and a reduction of root length (Fig. 5A and B). Thus, the loss of GLXI;2 activity in both loss-of-function plant lines impaired KDG detoxification. The complemented lines showed restoration of development, with root lengths similar to that of the wild type (Fig. 5A and B), demonstrating that the transgenic expression of GLXI;2 restores the metabolization of KDG in the loss-of-function seedlings.

**Figure 5.**
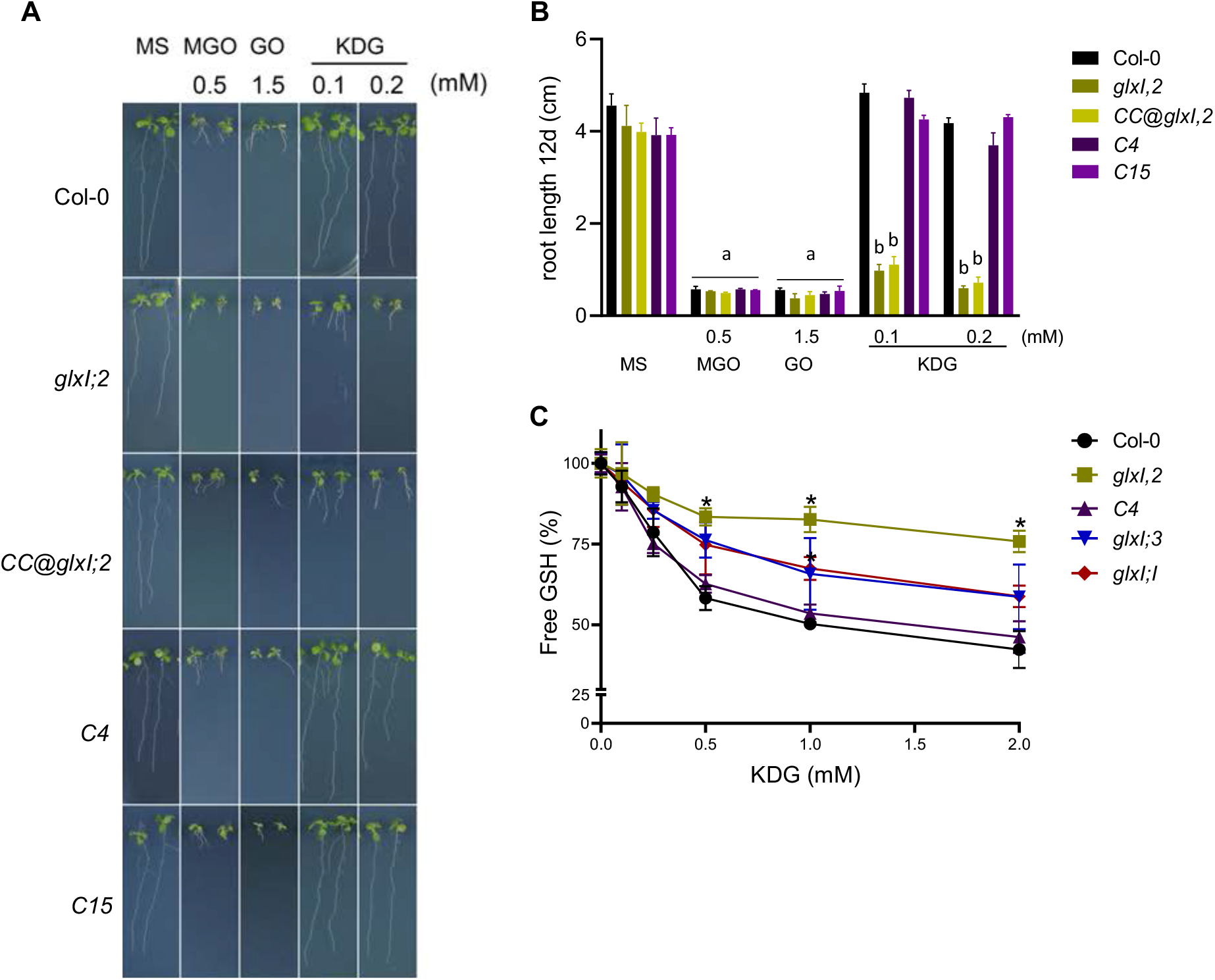
Effect of KDG on the seedling establishment of Arabidopsis GLXI loss-of-function and complemented lines. **A**, Representative picture of seedling development after 12 days of germination in MS medium and in the presence of MGO or KDG. Bar=1cm. **B**, Root length after 12 days of germination in MS medium and in the presence of MGO or KDG (as in A). Values are means ± SE from independent experimental trials (n=3-4) of 7-10 root measurements each. Testing for significant differences was performed by two-way ANOVA (ANOVA across treatments: P<0.0001; across genotypes: P<0.0001; interaction: P<0.0001). “a” and “b” indicate significant results of Tukey’s multiple comparison post-hoc test (P<0.001), where “a” indicates significant differences between treatment and MS control within a given genotype, and “b” indicates significant differences between genotype and Col-0 in a given treatment. **C**, Activity of GLXI using KDG as substrate in protein extracts of different plant lines. The free GSH concentration was determined after incubation of protein extracts of the wild type, loss-of function, and complemented lines with different KDG concentrations. 100% was considered the value of free GSH in mM when 0 mM of KDG was added. Values are means ± SD (n = 3). The asterisk (*) indicate significant differences between the values of the plant line and Col-0 at each KDG concentration (P<0.05), determined by Dunnett’s multiple comparisons test.

To directly demonstrate the action of GLXI;2 on KDG in plant tissues, we used protein extracts of wild-type, mutant, and complemented lines grown in standard conditions to perform GLXI activity assays. In the assays we used different concentrations of KDG and a fixed concentration of GSH. While KDG is metabolized by the wild-type protein extracts, it is hardly metabolized by protein extracts of the GLXI;2 loss-of-function mutant (Fig. 5C), consistent with the toxicity effects observed in the development of the loss-of-function mutants. Expression of GLXI;2 in the loss-of-function mutant background restored the ability of the protein extracts to metabolize KDG (Fig. 5C), consistent with the complementation of development of the seedlings in the presence of KDG. Protein extracts of *glxI;1* and *glxI;3* displayed similar rates of KDG consumption to the wild type, explaining the normal development of the seedlings in media containing KDG.

### 2-Keto-d-glucose affects root development of glxII;4 glxII;5 after germination

After the action of GLXI on a given RCS, the hemithioacetal produced is further metabolized through GLXII. Our phylogenetic analysis suggests that Arabidopsis GLXI;2, which represents the class unique to the viridiplantae, likely co-evolved with viridiplantae-specific GLXII;4 and GLXII;5 (Fig. 2 and Supplemental Fig 4). We thus speculate that GLXII;4 and/or GLXII;5 are mostly involved in the second step of KDG metabolization *in vivo*. To test this hypothesis, we produced homozygous loss-of-function mutants of all GLXII isoforms (Supplemental Fig. 8A-D). As GLXII;4 and GLXII;5 might have partially redundant functions in the metabolization of KDG, we also produced the double loss-of-function mutant *glxII;4-3/5-2* (from here on referred to as *glxII;4 5*) through genetic crosses. All the produced lines were used for feeding experiments with different RCS.

In the presence of MGO and GO, both GLXII;2 mutants presented smaller and more yellowish leaves than the wild type (Fig. 6A), while measurements of the root length indicated no significant differences (Fig. 6B). These results indicate that in the conditions of our assay, GLXII;2 most likely participates in the metabolization of the *S*-d-lactoylglutathione and S-glycolylglutathione produced by the action of GLXI;3 on MGO and GO, respectively. We also observed that the double mutant *glxII;4/5* was strongly affected by GO in the growth media, showing smaller and more yellowish leaves than the wild type (Fig. 6a), although the root length was similar to the wild type (Fig. 6B). These observations indicate that both viridiplantae-specific GLXII are also involved in the detoxification of *S*-glycolylglutathione and can complement each other. The loss-of-function lines of GLXII;4 and GLXII;5 were similar to the wild type in the presence of MGO and GO (Fig. 6A and B). When the seeds germinated in the presence of KDG, the double mutant *glxII;4/5* was highly affected in a dosedependent manner, showing a significant reduction of the root length after 12 days of grow (Fig. 6B). The loss-of-function mutant of GLXII;4 also showed a reduction of the root system when the higher concentration of KDG was used. These results indicate that both viridiplantae-specific GLXII are involved in the detoxification of the hemithioacetal formed by the metabolization of KDG via GLXI;2; in this process, GLXII;4 appears to have a dominant role.

**Figure 6.**
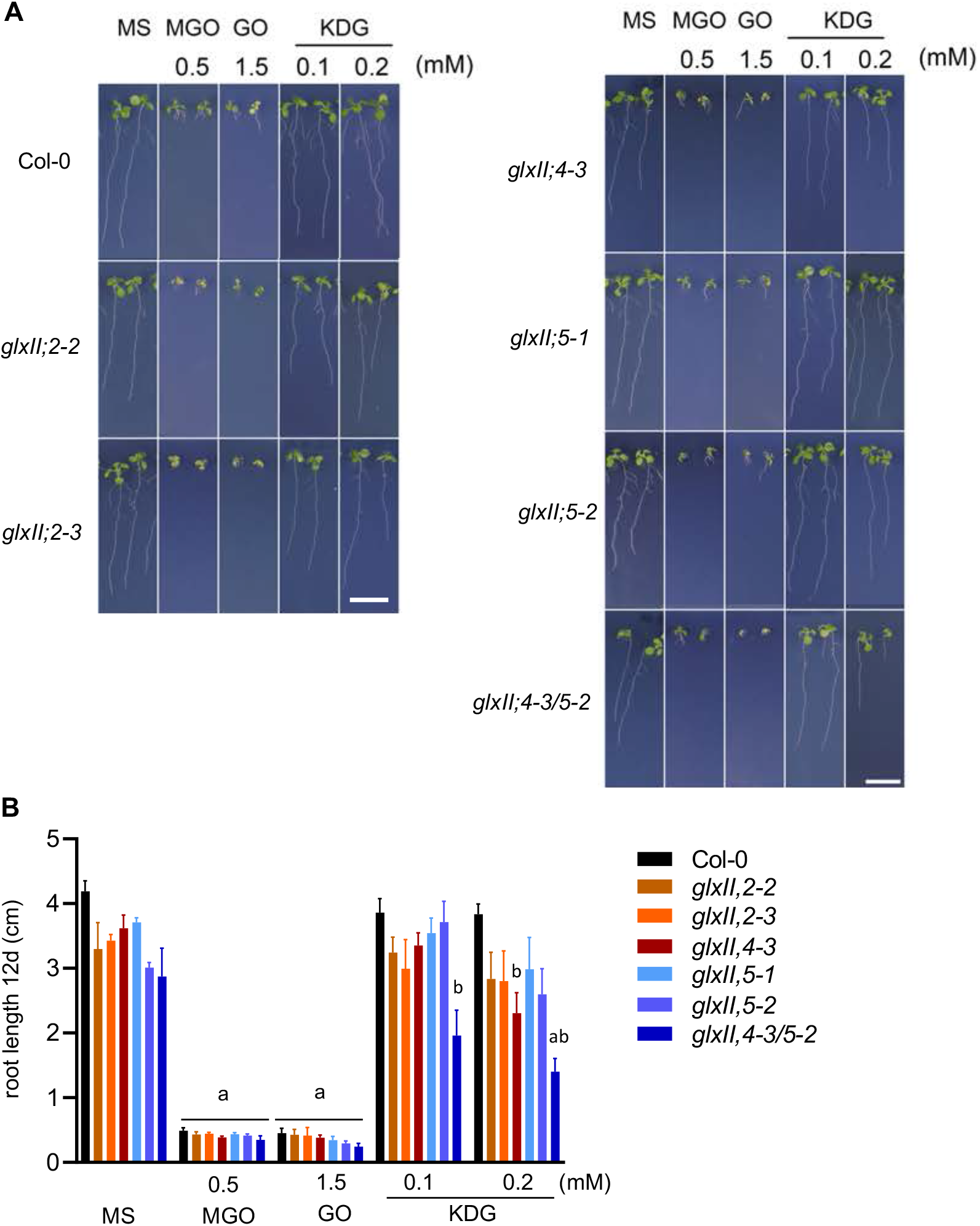
Effect of KDG, MGO and GO on the seedling establishment of Arabidopsis GLXII T-DNA insertion lines. **A**, Representative picture of seedling development after 12 days of germination in MS medium and in the presence of MGO, GO or KDG. Bar=1cm. **B**, Root length after 12 days of germination in MS medium and in the presence of MGO, GO or KDG (as in A). Values are means ± SE from independent experimental trials (n=3) with 7-10 root measurements each. Testing for significant differences was performed by two-way ANOVA (treatment: P<0.0001, genotype: P<0.0001, interaction: P<0.0001). “a” and “b” indicate significant results of Tukey’s multiple comparison post-hoc test (P<0.001), where “a” indicates significant differences between treatment and MS control within a given genotype, and “b” indicates significant differences between genotype and Col-0 in a given treatment.

### KDG levels vary in plant lines with modified expression of viridiplantae-specific GLXI and GLXII proteins

We speculated that if the wild type and GLXI;2 complemented lines are able to metabolize KDG present in the growth media, while the loss-of-function lines for GLXI;2 and GLXII;4/GLXII;5 are not able to detoxify it properly, these differences would be reflected in changes of the metabolic profiles of the plant tissues. We thus used GC-MS to conduct a metabolite profile analysis on root extracts of plants grown in the presence of 0.1 mM KDG. We performed a principal component analysis (PCA) of all samples, finding that principal component 2 clearly distinguished the two GLXI;2 loss-of-function mutants from all other lines analysed (Fig. 7A).

**Figure 7.**
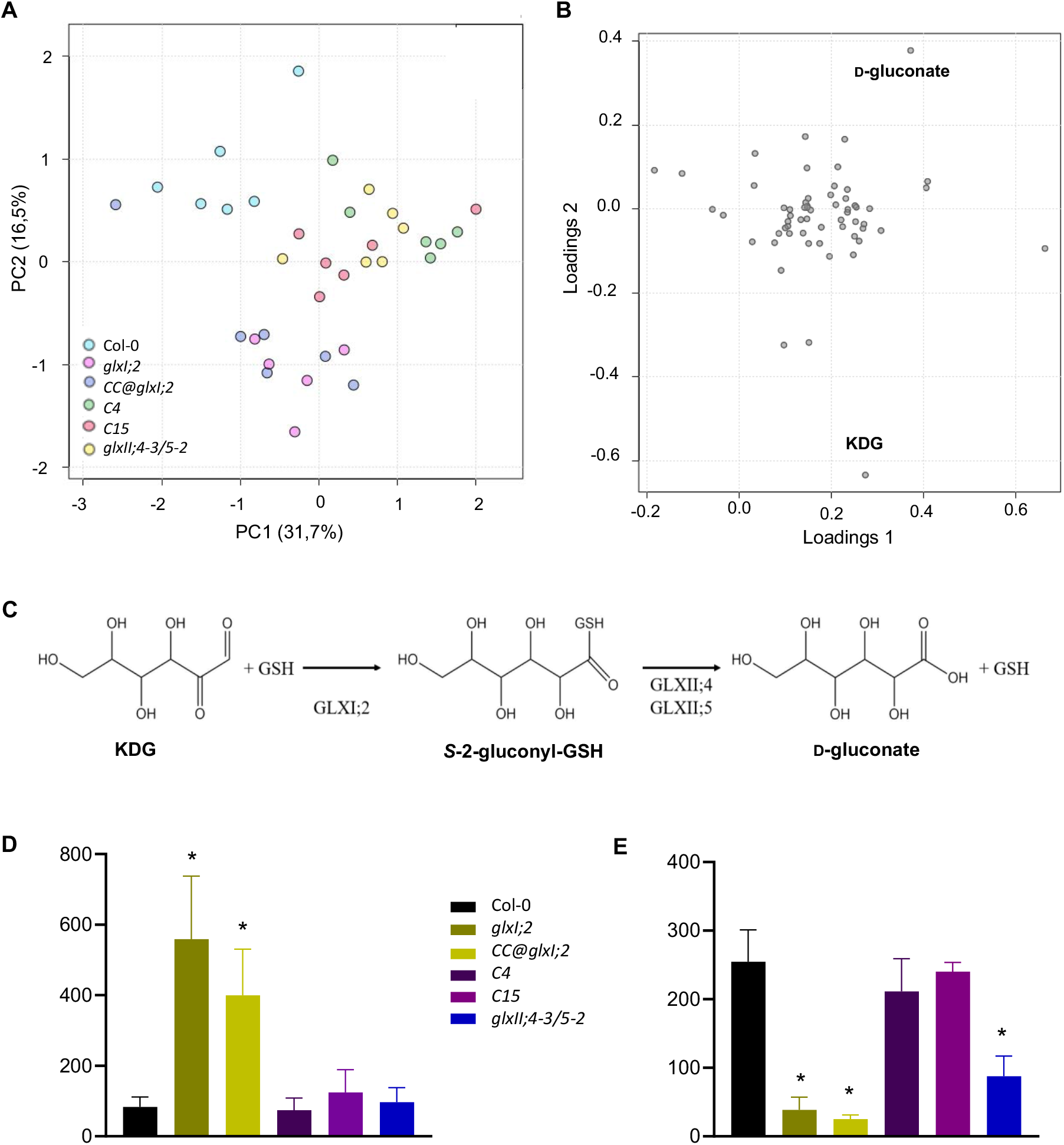
Metabolite analysis of roots of seedlings grown in the presence of KDG. **A**, Principal component analysis (PCA) of metabolites performed using the MetaboAnalyst 5.0 software. **B**, Loading plot of root metabolic profiles showing KDG and D-gluconate as the most discriminating metabolites. **C**, Depiction of the action of the GLX system on KDG. S-2-gluconyl-GSH is formed through the action of the GLXI;2 on KDG. This hemithioacetal is the substrate of GLXII;4/5, which produces d-gluconate. **D**, Relative amounts of KDG in Col-0, mutant and complemented lines quantified by GC-MS. **E**, Relative amounts of D-gluconate in Col-0, mutant, and complemented lines quantified by GC-MS. In D and E, bars and error bars show means and SD of six biological replicates. The asterisk (*) indicates significant differences between the mutant or complemented line and Col-0 (P<0.0001) as determined by the Dunnett’s multiple comparisons test.

A loading plot revealed that KDG and d-gluconate are the two metabolites that discriminate most strongly between the lines (Fig. 7B), consistent with KDG being the substrate of the viridiplantae-specific GLX system and d-gluconate being the corresponding product (Fig. 7C). We next zoomed in on the relative KDG levels in the plant lines. In the two mutant lines *glxI;2* and *CC@glxI;2*, KDG accumulated to much higher amounts than in the other lines (Fig. 7D), while the introduction of an active GLXI;2 into the *glxI;2* mutants reverted the accumulation of KDG to wild-type amounts (Fig. 7D). These results confirmed that in Arabidopsis roots, GLXI;2 metabolizes KDG. The roots of the GLXII double mutant *glxII;4/5* presented KDG levels comparable to the wild type, indicating the action of GLXI;2.

In the GLXI;2 mutant lines fed with KDG, we also expect a decrease of d-gluconate, the product resulting from the consumption of KDG by the plant-specific GLX system (Fig. 7c). The relative content of d-gluconate measured in roots of plants growing in KDG for 12 days indicated that in *glxI;2* and *CC@glxI;2* d-gluconate accumulates to only 15% and 10% of the levels seen in the wild type, respectively (Fig. 7E). The introduction of an active GLXI;2 into the *glxI;2* mutant returns d-gluconate to levels similar to the wild type (Fig. 7E). As expected, the double mutant *glxII;4/5* also presented lower relative levels of d-gluconate (34%) than the wild type (Fig. 7E). These results support our hypothesis that in Arabidopsis, KDG can be metabolized to d-gluconate through the consecutive action of GLXI;2 and GLXII;4 and GLXII;5.

### KDG produced from glucose impairs development of loss-of-function mutants of GLXI;2

As KDG is produced during cellular metabolism by the oxidation of glucose or the degradation of glycosylated proteins (Bean and Hassid, 1956; Baute, 1984; Semchyshyn, 2014), we hypothesized that plants growing in glucose produce enhanced levels of KDG. We confirmed that feeding with glucose increase the production of KDG in the plant tissues, finding that wildtype plants grown in the presence of 2% glucose presented 270% higher KDG amounts than those of seedlings grown in MS media (Fig. 8A). Furthermore, we analysed the development of plants with altered GLXI expression in glucose-enriched media. Glucose (2 %) in the growth media already impaired the development of the wild type, which showed smaller rosettes and a much shorter root system compared to growth in MS media (Fig. 8B). We found that *glxI;2* and *CC@glxI;2* development was much more affected than that of the wild type, presenting much smaller and yellowish rosettes. In contrast, the complemented lines behaved like the wild type (Fig. 8B). In the presence of glucose, the development of *glxI;3* was like the wild type, while the development of *glxI;1* was affected although to a much lesser extent (Fig. 8B). It is likely that GLXI;1 is not involved in the metabolization of KDG or that in the conditions of our analysis its activity on KDG is difficult to be assessed, as GLXI;1 is much less expressed in Arabidopsis tissues than GLXI;2 (Schmitz et al., 2017).

**Figure 8.**
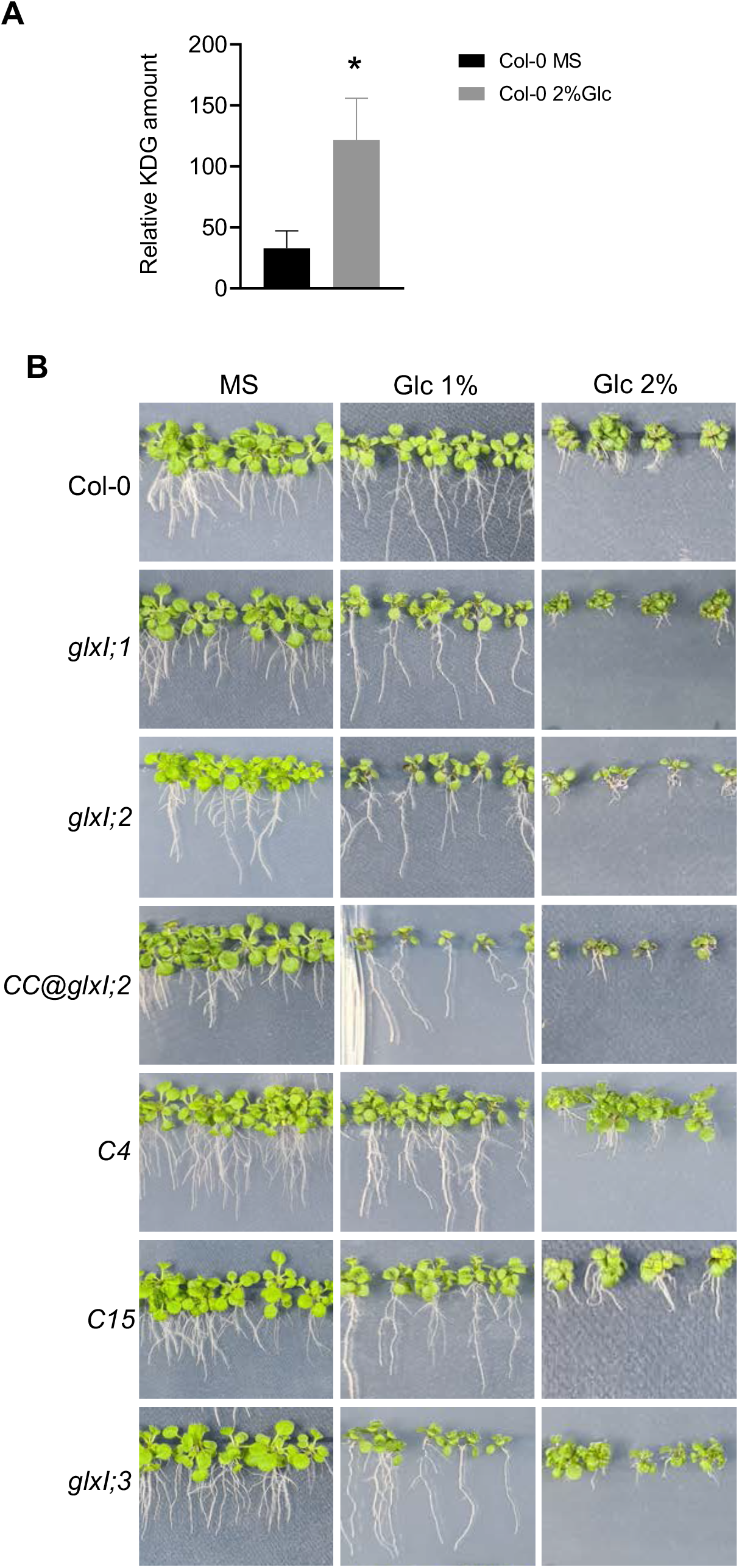
Effect of glucose on seedling development of Arabidopsis GLXI loss-of-function and complemented lines. **A,** Representative picture of seedling development after 12d of germination in MS medium and MS medium supplemented with 1 or 2% glucose (Glc). **B,** Relative amounts of KDG in Col-0 lines grown in MS and in 2% glucose quantified by GC-MS.

## DISCUSSION

In plants, members of the GLXI gene family form joint enzymatic pathways with members of the GLXII gene family. As indicated by the co-occurrence of the two gene families in genomes across bacteria and eukaryotes, the GLXI-GLXII cooperation likely already existed in ancient bacteria. All bacterial GLXI copies identified by our sequence searches show the conserved amino acids typical for Ni^2+^-preference, so it is likely that this represents the ancestral metal ion preference of GLXI. Our phylogenetic analyses identified GLXI and GLXII copies only in proteobacteria and in cyanobacteria, but not in other bacterial or archaeal clades. It appears likely that GLXI and GLXII genes were acquired by cyanobacteria through horizontal gene transfer from proteobacteria (Figs. 1 and 2). The most parsimonious scenario for the origin of the eukaryote-wide GLX system with Zn^2+^-type GLXI is that the endosymbiont of the primary endosymbiosis at the origin of eukaryotes, a proteobacterium, contributed genomic copies of GLXI and GLXII to the last common ancestor of all eukaryotes, LECA (Fig. 1B, bottom). These copies were inherited by all eukaryotic kingdoms, frequently involving gene duplications. As all GLXI enzymes of this monophyletic eukaryotic group show the conserved amino acids typical for Zn^2+^-preference, it is likely that this metal ion preference already existed in LECA. In all animals and plants analyzed, Zn^2+^-type GLXI possess one structural domain per monomer and are therefore active as dimers; in other eukaryotes studied, the proteins contain one or two structural domains. These domain duplications seem to have occurred independently in different lineages (Supplemental Fig. 1), representing an interesting case of convergent evolution.

Our phylogenetic analyses reveal a second, monophyletic set of GLXI and GLXII genes in viridiplantae. The corresponding gene copies have no close relatives among other eukaryotes, with the exception only of diatoms. The latter finding supports rather than contradicts the viridiplantae-specificity, as diatoms arose through secondary endosymbioses involving a green alga (Moustafa et al., 2009; Deschamps and Moreira, 2012). A potential, parsimonious explanation for the observed restriction of these GLX genes to viridiplantae would be that they came with the primary endosymbiont at the origin of photosynthetic eukaryotes. However, this endosymbiont was a cyanobacterium; in contrast, the closest relatives of both viridiplantae-specific GLXI and viridiplantae-specific GLXII genes are proteobacteria. This observation indicates that this GLX system was acquired by the lineage of the last common ancestor of viridiplantae through horizontal gene transfer from a proteobacterium. The GLXI genes of this clade have two domains, and the corresponding enzymes are active as monomers. Each domain forms a separate, monophyletic clade within the viridiplantae, indicating that the domain duplication must have occurred already in the last common ancestor of viridiplantae. In contrast, all bacterial (Ni^2+^-type) GLXI enzymes are dimeric, with one structural domain per monomer.

Why do viridiplantae have two GLX systems with different GLXI metal ion specificities? Existing literature on plant GLXI attributes the presence of plant GLXI isoforms to genetic redundancy and their biological function to the elimination of MGO (Kaur et al., 2013; Turra et al., 2015; Schmitz et al., 2017). Our results lead us to disagree with this view, for several reasons. In the monomeric Zn^2+^-type GLXI proteins, which exist in eukaryotes outside of plants and animals, the two structural domains feature conserved amino acids at the positions known to be involved in the interaction with the metal cofactor and with GSH. This suggests the presence of two functionally similar active sites, which were confirmed in the case of GLXI from *S. cerevisiae* (Frickel et al., 2001). In contrast, amino acid changes at the functional positions in one of the two domains in the viridiplantae-specific monomeric Ni^2+^-type GLXI (Fig. 3) lead to a modification of what is the second active site in the bacterial proteins. Structural analyses indicated that only sites in which the metal cofactor displays an octahedral coordination geometry are catalytically active (He et al., 2000; Suttisansanee et al., 2011 2015). Such an octahedral geometry arises from the metal coordination to four conserved residues present in His/Glu/His/Glu or His/Glu/Gln/Glu metal-binding motifs, completed by two solvating water molecules (Turra et al., 2015). As the conserved amino acids in the second domain of viridiplantae-specific GLXI differ from these two conformations, it appears likely that the corresponding monomeric enzymes possess only one active site, as has been suggested previously (Turra et al., 2015). The conservation of the corresponding amino acids strongly suggests that the modifications in Ni^2+^-type GLXI of viridiplantae are functionally important. This insight, together with the findings that the two-domain Ni^2+^-type isoforms are unique to viridiplantae and that the functional properties of this group differ from those of the Zn^2+^-type GLXI proteins, lead us to conclude that the Ni^2+^- and the Zn^2+^-type isoforms evolved in viridiplantae under distinct selective pressures. The domain phylogenies and the patterns of amino acid conservation indicate that the neofunctionalization of the two-domain Ni^2+^-type GLXI proteins must have occurred already in the last common ancestor of viridiplantae. We propose that GLXI;2 evolved in viridiplantae to scavenge KDG, an activity that is particularly important during seedling development. In contrast, the Zn^2+^-type GLXI;3 is crucial for the elimination of MGO and GO during both germination and seedling establishment (Schmitz et al., 2017).

In line with the general function of GLXI in glycation defense (Thornalley, 2003), we observed specific responses of the GLXI isoforms to alterations in cellular sugar levels (Fig. 8 and Ref. (Schmitz et al., 2017)). In contrast to our previous result that transcription of GLXI;2, and in some cases GLXI;1, is regulated by the cellular sugar status (Schmitz et al., 2017), we found that feeding *A. thaliana* with sucrose only affected the germination and seedling establishment of GLXI;3 loss-of-function mutants (Schmitz et al., 2017). As MGO is mostly formed from the activity of TPI during glycolysis (Richard, 1991; Phillips and Thornalley, 1993), feeding with sucrose most likely enhances glycolytic activities, thereby increasing the levels of MGO in the plant tissues. Interestingly, feeding with glucose impaired the development of GLXI;2 loss-of-function mutants in a similar way as KDG, and in plants grown in glucose-rich media, KDG accumulates to high levels. This observation suggests that KDG is produced through the oxidation of glucose or through the catabolism of glycosylated proteins formed by high glucose levels.

It is striking that viridiplantae are the only eukaryotes that possess both classes of GLX systems simultaneously. This uniqueness indicates that viridiplantae either suffer from more or more diverse stresses related to the production of RCS than other eukaryotes, or that other eukaryotes use complementary systems that compensate for their less diverse GLX systems. In mammals, two widely expressed enzyme families, aldehyde reductase and aldose reductase (EC1.1.1.21), are involved in the metabolization of 3-DG to 3-deoxyfructose and, to a lesser extent, of KDG to fructose, using NADPH as a cofactor (Kato et al., 1990; Takahashi et al., 1993; Feather et al., 1995). In contrast, there is no empirical evidence of the metabolization of KDG by homologs of aldehyde reductase and aldose reductase in the green lineage. Independently of the possible existence of such reductases in plants, we postulate that Ni^2+^-type GLXI type proteins in viridiplantae have evolved the ability to metabolize other alpha-oxoaldehydes in addition to MGO and GO, such as KDG, either as the main agents of detoxification or to accompany the activities of reductases in regulating cellular RCS levels. A complementary role of glyoxalases and reductases is suggested by the observation that aldose reductases, in addition to GLXI, are also very active on MGO in plants (Saito et al., 2013).

Consistent with the phylogenetic evidence that Ni^2+^-type GLXI proteins in viridiplantae coevolved with viridiplantae-specific GLXII, we found that both viridiplantae-specific GLXII genes in Arabidopsis, GLXII;4 and GLXII;5, are involved in the detoxification of the hemithioacetal formed by the metabolization of KDG through GLXI;2 (Fig. 6). GLXI;2 is found in the cytosol, while GLXII;4 and GLXII;5 are located in chloroplasts and mitochondria (Schmitz et al., 2017). The product of the GLXI;2 reaction, *S*-2-gluconyl-GSH, must thus be transported to the organelles. We hypothesize that GLXI;2 converts KGD present in the cytosol into *S*-2-gluconyl-GSH, which is then transported to the chloroplasts, where GLXII;4 and GLXII;5 produce d-gluconate for the chloroplastic localized pentose phosphate pathway (PPP), the main generator of cellular NADPH (Kruger and von Schaewen, 2003). The involvement of different subcellular compartments in a detoxification pathway is also observed during the metabolization of MGO. In this case, the last enzyme of the pathway, d-lactate dehydrogenase (Engqvist et al., 2009; Welchen et al., 2016), is found in the mitochondrial intermembrane space. Accordingly, either d-lactate produced in the chloroplast or cytosol by the sequential action of GLXI and GLXII must be transported to mitochondria (Schmitz et al., 2017), or the intermediate product of GLXI activity on MGO, S-lactoyl-GSH, is transported into the mitochondria to form –-lactate by the action of mitochondria-localized GLXII (Schmitz et al., 2017). The distribution of enzymatic activities of the GLX pathway across different organelles may optimize the provision of the corresponding precursors and/or the feeding of the final product into the relevant cellular metabolic pathways. This could be possible if there exist transient coupling of metabolizing enzymes with transporters to facilitate the transfer of substrates from one enzyme to the other sequential enzymes (Winkel, 2004; Sweetlove and Fernie, 2018).

The distinct properties of the GLXI isoforms and their co-existence in the same subcellular compartments (Schmitz et al., 2017) suggest that the isozymes perform specific metabolic functions in vivo. Our work shows that GLXI;2, a Ni^2+^-type GLXI type protein unique to viridiplantae, is involved in the detoxification of RCS other than MGO and GO, specifically of KDG. Importantly, we found that viridiplantae-specific Ni^2+^-type GLXI proteins co-evolved with plant-specific GLXII proteins. These results provide a strategy for improving plant resistance to RCS produced in response to abiotic stresses, by expressing pairs of GLX genes that work together to detoxify the relevant RCS (such as GLXI;2 and GLXII;4, or GLXI;3 and GLXII;2) in the respective cells.

## Methods

### Sequence selection, alignment and phylogenetic construction

Known GLXI and GLXII sequences from the five representative species *Homo sapiens, Arabidopsis thaliana, Phaeodactylum tricornutum, Synechocystis salina*, and *Escherichia coli* were selected as queries for a protein-protein blast (blastp) with an expectation value threshold of e<0.0005 against the RefSeq protein database (Pruitt et al., 2012). This search resulted in a total of 127 amino acid sequences for GLXI (Supplemental File 1) and in a total of 98 amino acid sequences for GLXII (Supplemental File 2).

We separately aligned all GLXI and all GLXII sequences, using the program Clustal Omega (Chojnacki et al., 2017) with default settings (Supplemental File 3 and 4). Some GLXI sequences from viridiplantae, rhodophyta, phaeophyta, diatoms, fungi, and protozoa comprise two copies of the structural domain found in the other GLXI sequences. We first aligned all two-domain sequences (Supplemental File 5) and then split them at the hinge region, identified from the 3-D protein structure (Turra et al., 2015), cutting the alignment at the position corresponding to E146 in *A. thaliana* GLXI;2 (Supplemental Fig 1). We denoted the resulting first and second domain sequences with the suffix “_A” and “_B”, respectively, and then aligned them as individual sequences with the one-domain GLXI sequences (Fig. 3C).

We inferred the maximum-likelihood trees for GLXI and GLXII alignments using the program MEGA X (Kumar et al., 2018). Initial trees for the heuristic tree topology searches were obtained using neighbor-joining, based on a matrix of pairwise distances estimated using the JTT model. For the subsequent maximum-likelihood analysis, we chose the LG model of sequence evolution (Le and Gascuel, 2008), which was the best-fitting model among those implemented in MEGA X for both alignments. A discrete Gamma distribution was used to model evolutionary rate differences among sites (5 categories). To estimate branch support values, we calculated trees for 1000 bootstrap replicates of each alignment.

Low bootstrap support values reflect uncertainties in the tree reconstruction. Such uncertainties are often caused by a small number of individual “rogue” sequences that cannot be placed reliably in the better-supported tree describing the relative positions of the remaining sequences. We identified such rogue taxa using the RogueNaRok algorithm and software (Aberer et al., 2013), following the methodology described in Russo et al. (Russo and Selvatti, 2018). Sequences with a majority rule consensus coefficient (*mr*) value (Aberer et al., 2013) above 0.5 were considered rogue and were eliminated. For GLXI, this strategy resulted in the removal of *Candidatus Pelagibacter* (*mr*=0.68), *Duganella phyllosphaerae* (*mr*=0.68), *Aliidongia dinghuensis (mr=0.57), Sphingomonas sanguinis (mr=0.56*), and *Magnetospirillum magneticum (mr=0.56*). From the GLXII alignment, the same strategy resulted in the removal of *Nitrospirillum amazonense (mr=0.63), Rhizomicrobium palustre (mr=0.62*), and *Sneathiella chinensis (mr=0.52*).

After removal of the rogue taxa from the FASTA files, we realigned the GLXI domains and the GLXII gene sequences, respectively. We then reconstructed the two trees again, as described above. Supplemental Fig. 2 and 4 show the trees with the highest likelihoods for GLXI and GLXII, respectively, together with the branch support values from 1000 bootstrap replicates.

To root each of the two trees, we first restricted the corresponding alignment to the bacterial sequences, as substitution rates can differ substantially between prokaryotes and eukaryotes (visible from the branch lengths in Supplemental Fig. 2 and 4). For each reduced alignment, we then reconstructed the bacterial gene family tree as described above. We identified the root of this bacterial tree using the *minimal ancestor deviation* (MAD) algorithm (Tria et al., 2017), which minimizes the average distance between the root and all tips of the tree. We then used FigTree v1.4.4 (http://tree.bio.ed.ac.uk/software/figtree/) to display and root the complete tree containing all bacterial and eukaryotic sequences at the corresponding branch (Supplemental Fig. 2 and 4).

For the main text Figures 1 and 2, we summarized the full trees for GLXI (Supplemental Fig. 2) and GLXII (Supplemental Fig. 4) as follows. (i) We collapsed all branches with bootstrap support values below 50% to polytomies, using the program TreeCollapserCL4 (http://emmahodcroft.com/TreeCollapseCL3.htmlS), as the splits corresponding to these branches are deemed unreliable. (ii) For better readability, we grouped monophyletic sets of sequences that belonged to the same clade of organisms into sequence clusters (represented by triangles in the figures). (iii) Additionally, for GLXI and GLXII, we merged three and five bacterial sequences, respectively, into monophyletic bacterial clusters that descended from the same polytomy resulting from step i, identifiable in the Figures by the missing bootstrap support values at the base of the corresponding triangles.

### Plant lines and growth conditions

*Arabidopsis thaliana* Ecotype Columbia-0 (Col-0) was used. *A. thaliana* T-DNA insertion lines in *GLXII* were obtained from The European Arabidopsis Stock Centre and screened for loss-of function on genomic and transcriptional level using specific primers (listed in the T-DNA screening section in Supplemental Table 1). We identified homozygous insertion lines for all *GLXII*; for *GLXII;2*, lines *glxII;2-2* (SALK_090826) and *glxII;2-3* (SALK_134008); for *GLXII;4*, line *glxII;4-3* (SALK_045876); and for *GLXII;5*, lines *glxII;5-1* (SALK_014157) and *glxII;5-2* (SALK_100618) (Supplemental Fig. 8). Remaining transcript levels of GLXII in the insertion lines were analysed by RT-PCR as described below and using specific primers (listed in the RT-PCR for GLXII T-DNA insertion lines section in Supplemental Table 1). The *glxII;2-2* and *glxII;2-3* lines have relative residual transcript levels of 8.8% and 12.4%, respectively, based on band intensity quantification relative to the wild type using Fiji (Schindelin et al., 2012). We produced lines lacking the organellar GLXII activities by genetic crossed of *glxII;4-3* with *glxII;5-2*. Arabidopsis T-DNA insertion lines in *GLXI* were obtained in our previous work (Schmitz et al., 2017).

All plant lines were grown, on soil or sterile media, at 22°C under a long-day (16 h light/8 h darkness) photoperiod using a mix of bulbs from Osram HO 80W/840 and HO 80W/830 with a light intensity of 120 μE·m^−2^·s^−1^. Seeds grown on solid sterile culture (0.5x MS medium solidified with 0.9 % plant agar (Duchefa; Murashige and Skoog (1962)) were surface sterilized with ethanol according to standard protocols and individually transferred on plates containing MS only or supplemented with either 0.5 mM MGO, 1.5 mM GO, 0.1 or 0.2 mM KDG, 0.2 3DG, 2% or 4% glucose. All these metabolites were added after autoclaving to avoid oxidation. Seedson soil or plates were stratified 2 d at 4 °C prior to germination in standard growth conditions. Root length was documented photographically after 12 d of germination and analysed with ImageJ software (Schindelin et al., 2012).

### Complementation of glxI;2

Full length genomic sequence of GLXI;2 was used to complement the *glxI;2* T-DNA insertion line. Arabidopsis Col-0 genomic DNA was used as template to amplify the sequence using Phusion Polymerase (Thermo Scientific) and the specific primers OX-GLXI;2_F and OX-GLXI;2_R (Supplemental Table 1). The PCR fragment obtained was subcloned into p-TOPO-Blunt, sequenced and subsequently amplified for vector assembly according to (Gibson, 2009) using the specific primers pHyg_F and MS_gly1-2_R (Supplemental Table 1). The vector pUBQ10_Venus was linearized with SacI and XhoI to remove the Venus tag and assembled with the GLXI;2 gene fragment to express GLXI;2 constitutively under the control of the AtUBQ10 promoter. The vector pUBQ10-gGLXI;2 was transformed into *Agrobacterium tumefaciens* (strain GV3101 pMP90) and selected on YEB plates containing rifampicin (150 μg/mL), gentamycin (50 μg/mL), and kanamycin (50 μg/mL). For stable expression in Arabidopsis, *glxI;2* T-DNA insertion plants (Schmitz et al., 2017) were transformed according to standard protocols and positive transformants were selected on 0.5x MS and hygromycin (50 μg/mL). Successful complementation of *glxI;2* by pUBQ10-gGLXI;2 was confirmed in three independent complementation lines (C4, C10, and C15) by amplification of genomic DNA and also at the transcriptional level.

### Generation of an independent glxI;2 mutant line using the CRISPR/Cas9 system

The CRISPR/Cas9 system was used to generate an independent loss-of-function line for the GLXI;2 gene locus. Small guide RNA (sgRNA) design, vector construction and positive selection was performed as described in Hahn, et al. (Hahn et al., 2017). In brief, sgRNAs were designed using CHOPCHOP (Labun et al., 2016) and two sgRNAs were selected targeting exon 2 or/and exon 4 (Supplemental Fig. 6a). The sgRNAs were generated with the specific primers pFH6_1_GLXI;2 and pFH6_2_GLXI;2 listed in Supplemental Table 1 in the GLXI;2 sgRNA section and subcloned into pFH6_new resulting in the plasmids pFH6_new-sgRNA1 and pFH_new-sgRNA2. The plasmids were used as templates for construction of fragments to be assembled with pUB-Cas9 vector according to Gibson (2009). The resulting vector pUB-Cas9@glxI;2 (sgRNA1+2) was transformed via *A. tumefaciens* (strain GV3101 pMP90) into Arabidopsis Col-0 according to standard protocols. Positive hygromycin transformants (T1) were screened via PCR for the presence of the pUB-Cas9@glxI;2 construct and for activity of Cas9 through PCR and restriction fragment length polymorphism (Supplemental Fig. 6a-e). The primers used are shown in Supplemental Figure 6 and listed in Supplemental Table 1 in the CC@glxI;2 screening section. The T2 generation of selected plant lines (CC@glxI;2-1, −3, −6, and −8) were screened for loss of pUB-Cas9@glxI;2 and stable mutation of sgRNA target sites via PCR, and PCR fragments were sequenced to confirm the homozygous mutation at the target site. For further characterization we selected plants of the T3 generation of line CC@glxI;2-1, as we confirmed a homozygous deletion between exons 2 and 4 in these plants (Supplemental Fig. 6A).

### RNA isolation from plant tissue and synthesis of complementary DNA

RNA was isolated from rosette leaves using the Universal RNA Purification Kit from Roboklon GmbH (Germany) according to the protocol provided by the manufacturer. Isolated RNA was eluted in 30 μl RNase-free dH_2_O. The purity of the isolated RNA was checked using the photospectrometer NanoDrop ND 1000 (PEQLAB Biotechnologie GmbH, Germany) and subsequent agarose gel electrophoresis. The total RNA extracted was kept at −80 °C until use.

The synthesis of cDNA from 2 μg of RNA was carried out in two steps. In the first step, DNA was removed from the samples using DNA-free kit (Life Technologies Inc, USA) following the manufacturer’s guidelines. In the second step, cDNA was synthesized from RNA by reverse transcription. One μl of oligo(dT) primer was added to the RNA and incubated at 65° C for 10 min. Subsequently, 2 μl of dNTPs, 0.8 μl RevertAid H Minus M-MuLV Reverse Transcriptase (ThermoFischer, USA), 4 μl of 5X RevertAid H Minus First Strand cDNA Synthesis Kit reaction buffer (ThermoFischer, USA) and H_2_O up to 10 μl, was added to the RNA. The mixture was incubated at 42° C for 60 min and then inactivated at 70° C for 10 min. The cDNA was stored at −20° C until use.

### Semiquantitative PCR (RT-PCR) and Quantitative (qPCR)

RT-PCR was performed with cDNA extracted from plants of line *CC@glxI;2-1*, and the primer pair Gly-h-At1g11840 FW + Gly-h-At1g11840 RV (section RT-PCR for CC@glxI;2 in Supplemental Table 1). The expected PCR product size was 852 bp in wild-type plants and 686 bp in plants with deletion. The housekeeping gene ACTIN2 (Czechowski et al., 2005), amplified with the primers Actin_F+ Actin_R (section RT-PCR for T-DNA insertion lines of Supplemental Table 1), served as the internal standard. Agarose (1%) gel electrophoresis was used to evaluate the size of all PCR products shown in Supplemental Figure 6A.

Transcript analysis of *GLXI;2* in the complemented lines C4, C10, and C15 was conducted via qPCR using KAPA SYBR FAST qPCR Master Mix (Kapa Biosystems, Roche, Switzerland) and 2 mL of 1:10 diluted cDNA in an Applied Biosystems Step One Plus real-time PCR instrument (ThermoFisher, USA), according to the manufacturer’s instructions. ACTIN2 (ACT2) was used as reference gene (Czechowski et al., 2005). The primers used to amplify the *GLXI;2* transcript, RL_GLXI;2_F and RL_GLXI;2_R, and ACT2, RL_Act_F* and RL_Act_R* (section GLXI;2 qPCR of Supplemental Table 1) were produced using Primer-Blast (Ye et al., 2012). Differential *GLXI;2* gene expression was analyzed and expressed using the ΔΔCT method (Schmittgen and Livak, 2008).

### Protein extraction and GLXI enzymatic assay

The plant lines were germinated and grown 14 d in plates and whole plantlets were harvested in the middle of the day. For total protein extraction 200 mg of plant material was grinded in the presence of liquid N2 and extraction buffer (100 mM Tris-HCl, pH 8.0; 100 mM NaCl; 0.5 % (v/v) Triton X-100; 2 mM PMSF; 1 % (w/v) PVP40). The samples were centrifugated at 4°C for 10 min at 20000g, and the supernatant was transferred to a clean tube.

The enzymatic activity of GLXI was analysed by measuring free GSH in the media after completion of the reaction. Before starting the assay, KDG (2-Keto-D-glucose; Sigma-Aldrich, USA) was incubated with GSH for 10 min to enable the formation of the hemithioacetal. The concentration of GSH was kept at 1 mM, while variable KDG concentrations were used (0, 0.25, 0.5, 1 and 2 mM). Afterwards, 20 ug of protein extract was added to the reaction mixture and the absorbance was measured at 412 nm after 30 minutes of incubation. Finally, 1 mM DNTB (5, 5-dithio-bis (2-nitrobenzoic acid); Geno Technologies Inc., USA), which reacts with GSH, was added to the assay and the absorbance was measured again at 412 nm after 10 min of incubation. All assays were carried out with a Synergy HT plate reader (Biotek Instruments Inc., USA) at 25° C. All reagents of the assay were dissolved in 0.1 M MOPS pH 7.0. The determination of GSH concentration were calculated with a calibration curve performed with 0.2, 0.3, 0.4, 0.6, 0.8, 1.2, and 2 mM of GSH in 0.1 M MOPS pH 7.0. All assays were performed on at least three biological replicates each in at least triplicate determinations.

### Gas chromatography–mass spectrometry analysis

Root samples were harvested in the middle of the day/light after gown for 14 d in 0.5X MS or 0.1 mM KDG. Extraction and analysis by gas chromatography–mass spectrometry (GC-MS) were performed using the same equipment set-up and same protocol as described in Lisec et al. (2006). Briefly, 10 mg frozen material was homogenized in 300 μL of methanol at 70 °C for 15 min and 200 μL of chloroform, followed by addition of 300 μL of water. The polar fraction was dried under vacuum, and the residue was derivatized for 120 min at 37 °C (in 40 μl of 20 mg ml^−1^ methoxyamine hydrochloride (Sigma-Aldrich, cat. no. 593-56-6) in pyridine followed by a 30 min treatment at 37 °C with 70 μl of N-methyl-N-(trimethylsilyl)trifluoroacetamide (MSTFA reagent; Macherey-Nagel, cat. no. 24589-78-4). An autosampler Gerstel Multi-Purpose system (Gerstel GmbH & Co.KG, Mülheim an der Ruhr, Germany) was used to inject 1μl of the samples in splitless mode to a chromatograph coupled to a time-of-flight mass spectrometer system (Leco Pegasus HT TOF-MS; Leco Corp., St Joseph, MI, USA). Helium was used as carrier gas at a constant flow rate of 2 ml s-1 and GC was performed on a 30 m DB-35 column (capillary column, 30 m length, 0.32 mm inner diameter, 0.25 μm film thickness, PN: G42, Agilent). The injection temperature was 230 °C and the transfer line and ion source were set to 250 °C. The initial temperature of the oven (85 °C) increased at a rate of 15 °C min^−1^ up to a final temperature of 360 °C. After a solvent delay of 180 s, mass spectra were recorded at 20 scans s^−1^ with m/z 70–600 scanning range. Chromatograms and mass spectra were evaluated using Chroma TOF 4.5 (Leco) and TagFinder 4.2 software. Metabolites were annotated based on a retention index calculation with deviation <5% and compared with the reference data of the Golm Metabolome Database (http://gmd.mpimp-golm.mpg.de; (Luedemann et al., 2008)). The results were statistically analysed using Student’s *t*-test, using a threshold of P<0.05 between the samples of the control and treated plants (Supplemental File 6).

### Statistical analysis

P values were calculated using the Student’s t-test, one-way ANOVA with Dunnett test, or twoway ANOVA with Tukey’s test depending on the analysis performed. In each figure we clarify the analysis performed and the P values. A summary of the statistical results is presented in Supplemental File 7. Metabolic data were analyzed using metaboanalyst (https://www.metaboanalyst.ca/; (Pang et al., 2021)).

### Accession Numbers

Sequence data from this article can be found in the Arabidopsis Genome Initiative or GenBank/EMBL libraries under the following accession numbers: GLXI;1 (At1g67280), GLXI;2 (At1g11840), GLXI;3 (At1g08110), AtGLXII;2 (At3g10850), AtGLXII;4 (At1g06130) and AtGLXII;5 (At2g31350), HsGLXI (IAAV38791), EcGLXI (BAE76494), SsGLXI (TIGR00068), CvGLXI;1 (CHLNCDRAFT_48679), CvGLXI;2 (CHLNCDRAFT_134426), PtGLXI (PHATRDRAFT_12663).

## Supporting information

Supplemental Dataset6_GC-MS_Analysis

Supplemental Dataset7. Statistic analysis

## SUPPLEMENTAL DATA

**Supplemental Text.** Review of Kaur et al. 2013

**Supplemental Figure 1.** 3D structure of *A. thaliana* GLXI;2 obtained using the Phyre2 server prediction (Kelley, 2015) showing the typical fold defining the GLXI family.

**Supplemental Figure 2.** GLXI evolutionary tree, representing each domain of two-domain sequences as an individual sequence.

**Supplemental Figure 3.** Alignment of the viridiplantae GLXI proteins as shown in Fig. 2 and Supplemental Fig. 1.

**Supplemental Figure 4.** Evolutionary tree of GLXII sequences.

**Supplemental Figure 5.** Alignment of of viridiplantae GLXII proteins belonging to the *A. thaliana* GLXII;4 and GLXII;5 cluster in Fig. 3 and Supplemental Fig. 2.

**Supplemental Figure 6.** Arabidopsis GLXI;2 CRISPR/Cas9 generated loss-of-function lines.

**Supplemental Figure 7.** Analysis of *GLXI;2* expression in Arabidopsis *GLXI;2* loss of function and complemented lines.

**Supplemental Figure 8.** Arabidopsis GLXII T-DNA insertion lines.

**Supplemental Table 1.** Sequence of the primers used in this study.

**Supplemental Dataset 1.** Amino acid sequences corresponding to 127 GLXI proteins obtained from blastp.

**Supplemental Dataset 2.** Amino acid sequences corresponding to 98 GLXII obtained from blastp.

**Supplemental Dataset 3.** Alignment of the 127 amino acid sequences corresponding to GLXI by Clustal Omega.

**Supplemental Dataset 4.** Alignment of the 98 amino acid sequences corresponding to GLXII by Clustal Omega.

**Supplemental Dataset 5.** Alignment of the 34 amino acid sequences corresponding to the two-domain GLXI by Clustal Omega.

**Supplemental Dataset 6.** Table of relative metabolite amounts as determined by GC-MS. In each root line, there are six biological replicates (R1-R6).

**Supplemental Dataset 7.** Summary of statistical analyses.

## AKNOWLEDGEMENTS

This work was supported by the Deutsche Forschungsgemeinschaft through grant MA2379/21-1 to VGM; and, the Germany’s Excellence Strategy, through EXC2048/1 (Project ID:390686111) to MJL; and by the European Union’s Horizon 2020 research and innovation program, under PlantaSYST (SGA-CSA no. 739582 under FPA no. 664620) to SA and ARF.

The authors have no competing interests (financial/non-financial) that might be perceived to influence the interpretation of the article.

## AUTHOR CONTRIBUTIONS

JS and VGM conceived and designed research, and supervised data analysis and interpretation. MB, JS, MD, ICW, MS, SA performed research and acquired the raw data. ARF contributed with analytical tools. MB, JS, MJL and VGM wrote the manuscript. All authors contributed to the generation of the figures and the revision of the manuscript. All authors accepted the final version of the manuscript.

