## Supplementary material for "Viridiplantae-specific GLXI and GLXII isoforms co-evolved and detoxify glucosone *in planta*": Suppl Text, Figures and Table

**Supplementary Text: Review of Kaur et al. 2013**

In a publication entitled "Episodes of horizontal gene-transfer and gene-fusion led to co-existence of different metal-ion specific glyoxalase I" {Kaur, 2013}, Kaur et al. explored a subset of the questions addressed in our paper. Unfortunately, this earlier work could not provide reliable answers, as outlined in the following.

A major methodological problem is that it is unclear how exactly the authors assigned GLXI homologs to either the $Ni^{2+}$- or the $Zn^{2+}$-dependent GLXI type. The legend of their Supplementary Fig. S1 suggests that the presence of two sequence insertions was used as a distinguishing feature. However, this is neither explained nor justified in the text. The assignment appears to be in part based on sequence similarity to a GLXI homolog in *P. aeruginosa*, based on their Ref. 16; however, the latter publication could not reliably identify the corresponding metal ion specificity. Furthermore, while the Methods section reports that two rice proteins were "checked for the metal ion activation", the results are not shown; instead, the text refers to "our unpublished work" (p.9, Methods: "Assessment of metal ion specificity of GlxI"). The authors make several strong claims about the presence/absence and the evolution of the two forms with $Ni^{2+}$- and $Zn^{2+}$-dependence, as also indicated by their title. However, given that it is not clear how GLXI homologs were assigned to the two types, these claims cannot be justified with the evidence presented.

The following bullet points quote the main conclusions of Ref. {Kaur, 2013} and discuss their support.

- $Ni^{2+}$-dependent GLXI "evolved into Zn activated GlxI [Zn-GlxI] in deltaproteobacteria." – The authors' conclusion that $Zn^{2+}$-dependent GLXI exists in deltaproteobacterial genomes is based on sequence similarity to a GLXI homolog in *P. aeruginosa*. However, the metal ion specificity of that homolog is unclear, and hence the conclusion appears to be unwarranted.
- The "origin of eukaryotic Zn-GlxI is different and can be traced to GlxI from *Candidatus pelagibacter* and *Sphingomonas*." – This statement appears to be a misinterpretation of the domain tree in Fig. 5 of Ref. {Kaur, 2013}. The tree shows *Candidatus pelagibacter* as part of a larger bacterial sister group to the eukaryotic $Zn^{2+}$-dependent GLXI type sequences. *Sphingomonas* GLXI is the only bacterial sequence in the otherwise monophyletic eukaryotic group. However, it branches not basal to the eukaryotic sequences, but within the viridiplantae. Thus, it is unlikely that its lineage donated GLXI to eukaryotes. The tree topology instead is most consistent with either a sequence mis-annotation, a tree reconstruction artifact, or an (unlikely) horizontal gene transfer from algae to *Sphingomonas*.
- "In eukaryotes GlxI has evolved as two-domain protein but the corresponding Zn form is lost in plants/higher eukaryotes." – This statement is also not supported by the data presented. If the domain duplication had already occurred in a eukaryotic common ancestor, then the two domains should form separate clades that branch at the base of eukaryotes. Instead, the domains split within the apicomplexa/diatoms/brown algae clade, indicating an origin of two-domain $Zn^{2+}$-dependent GLXI in this group.

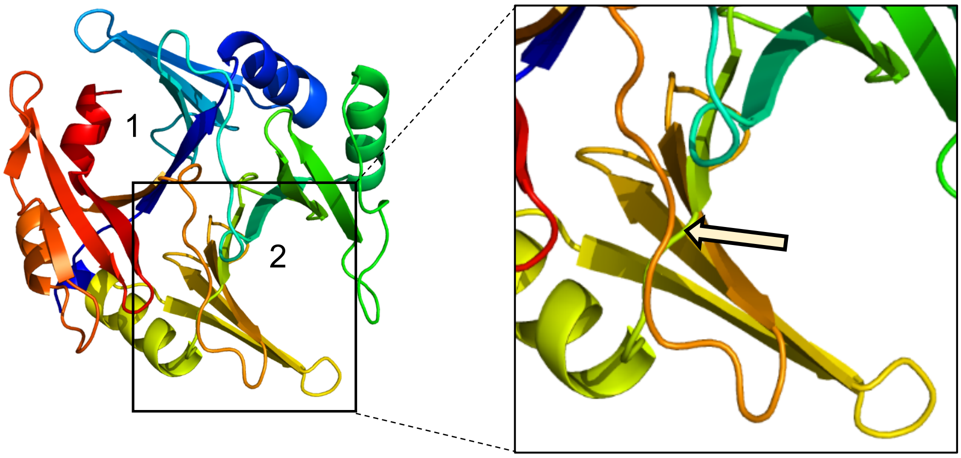


**Supplemental Figure 1. 3D structure of *A. thaliana* GLXI;2 obtained using the Phyre2 server prediction (Kelley, 2015) showing the typical fold defining the GLXI family.** The folding of the proteins allows the formation of two similar active sites (1 and 2), with participation of amino acids of both structural domains. To the right, zoom in of the hinge region (between the yellow and green colored regions that formed the active site 2), where the yellow arrow indicates the position corresponding to E146 that splits the protein into structural domains A (red and yellow regions) and B (green and blue regions) (Turra et al. 2015).

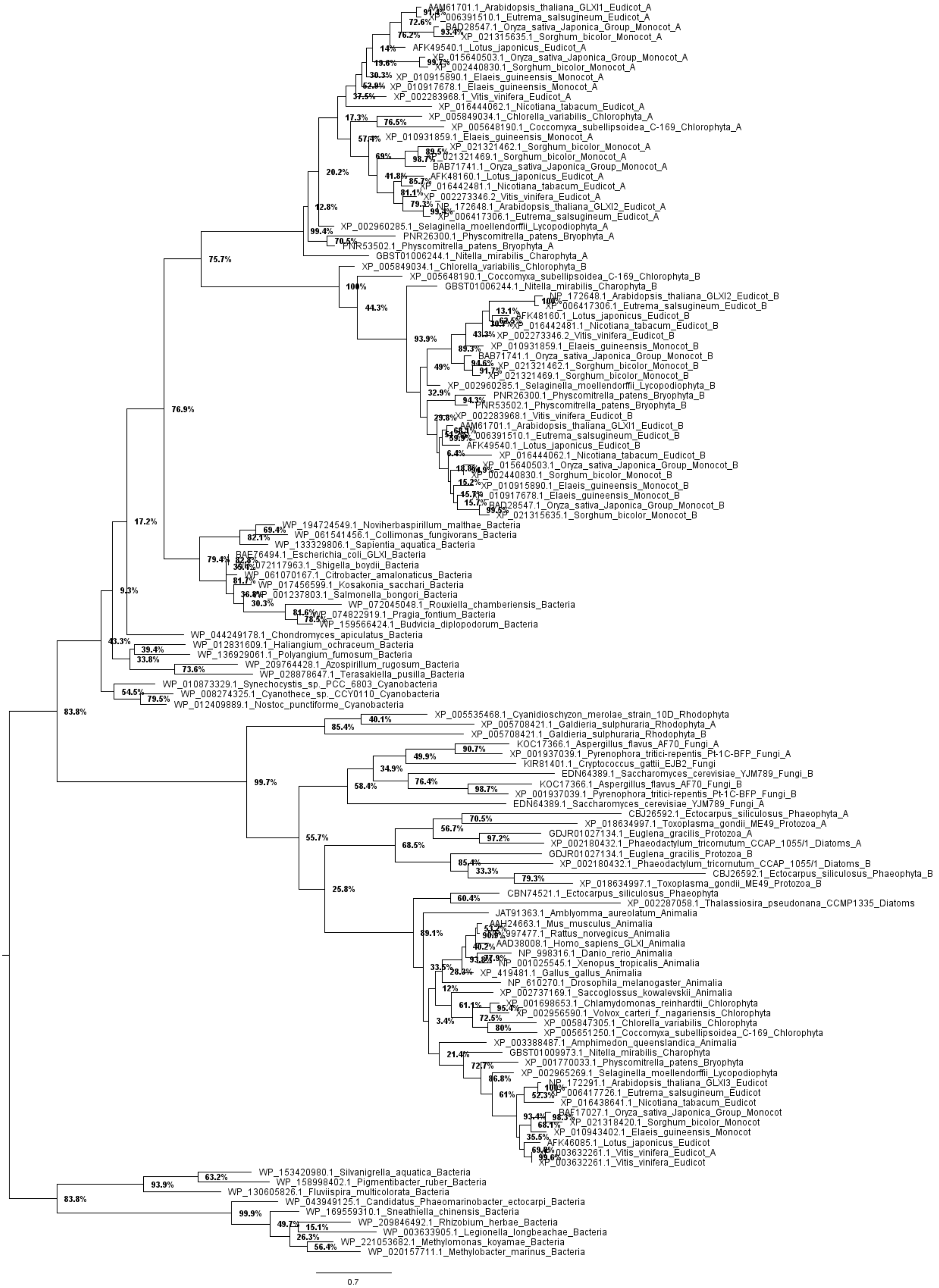


**Supplemental Figure 2. GLXI evolutionary tree, representing each domain of two-domain sequences as an individual sequence.** The figure shows the majority-rule consensus tree from 1000 bootstrap replicates; five rogue taxa were removed as detailed in Methods. Numbers atop branches indicate bootstrap support values in percent. A summary of this tree is shown in Fig. 1.

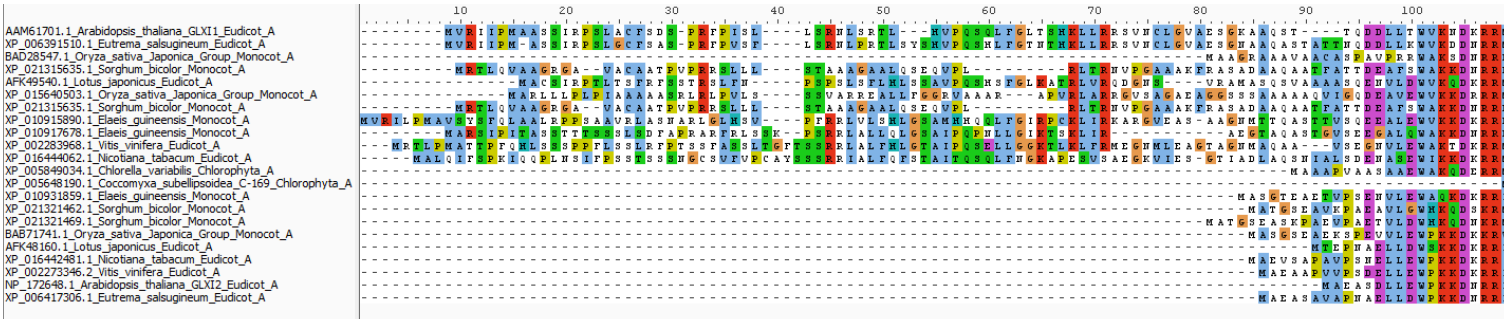


**Supplemental Figure 3. Alignment of the viridiplantae GLXI proteins as shown in Fig. 2 and Supplemental Fig. 1.**

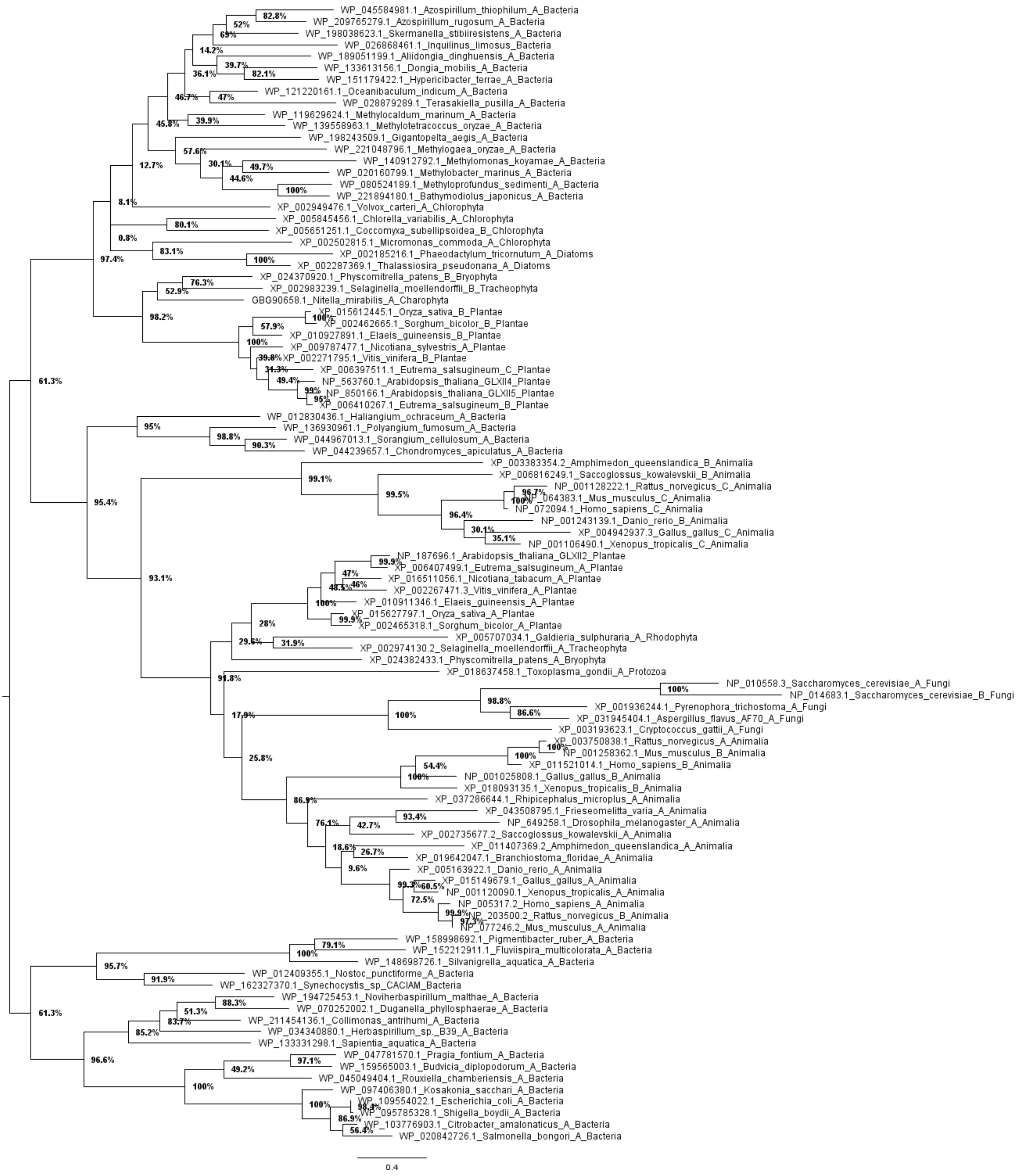


**Supplemental Figure 4. Evolutionary tree of GLXII sequences.** The figure shows the majority-rule consensus tree from 1000 bootstrap replicates; three rogue taxa were removed as detailed in Methods. Numbers atop branches indicate bootstrap support values in percent. A summary of this tree is shown in Fig. 3.

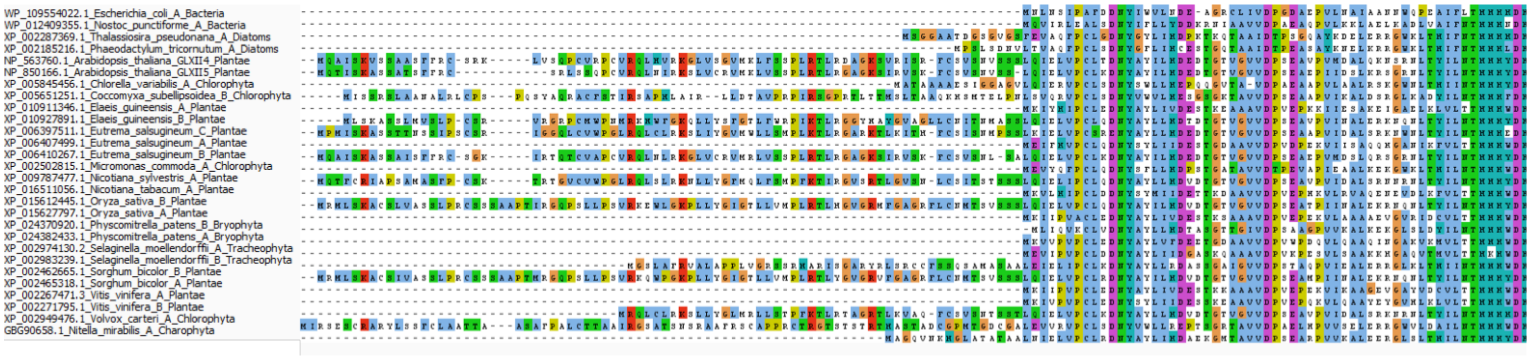

**Supplemental Figure 5. Alignment of of viridiplantae GLXII proteins belonging to the *A. thaliana* GLXII;4 and GLXII;5 cluster in Fig. 3 and Supplemental Fig. 2.**

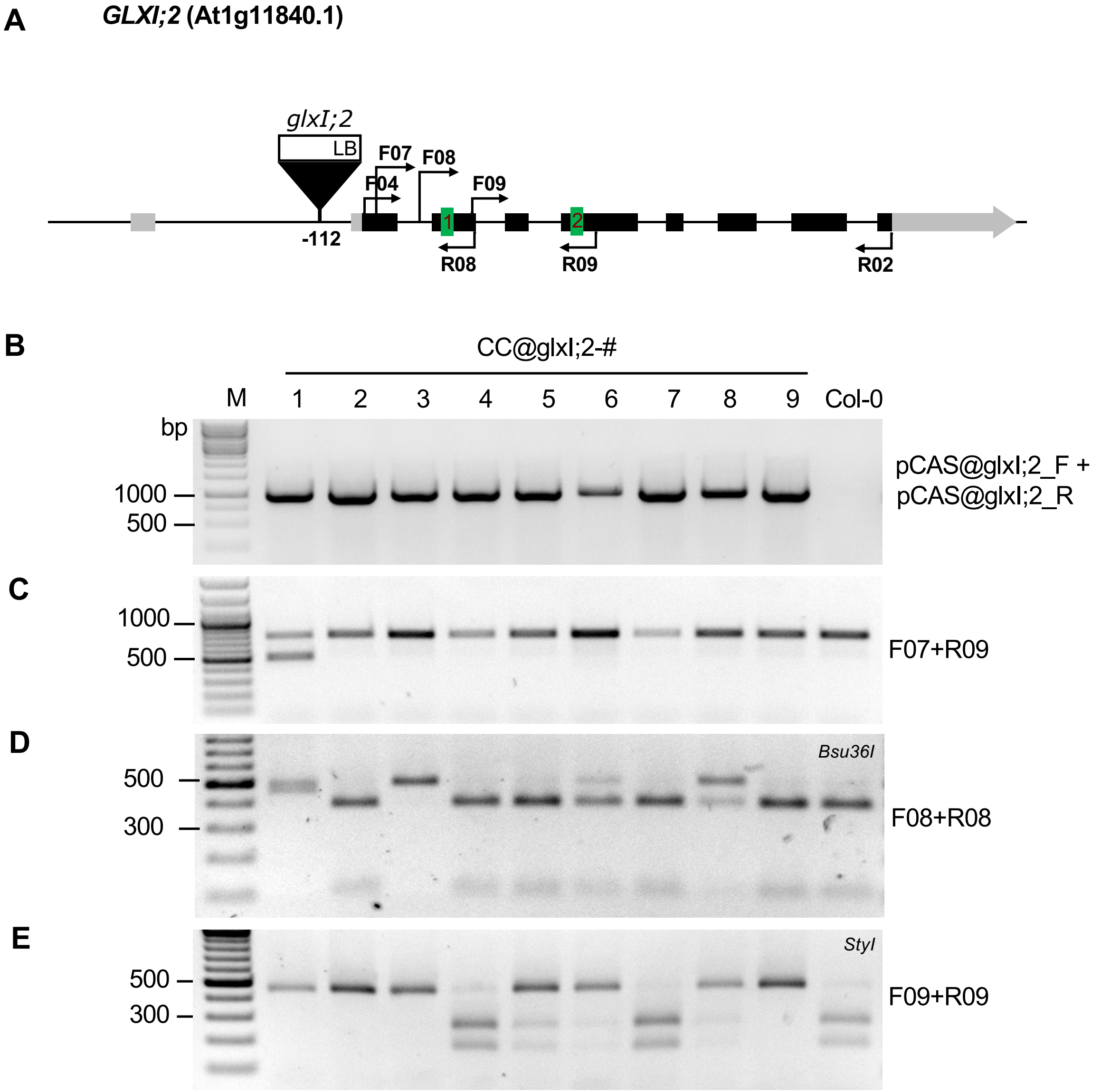


**Supplemental Figure 6. Arabidopsis GLXI;2 CRISPR/Cas9 generated loss-of-function lines.** **A,** Schematic representation of the *GLXI;2* gene locus showing the verified T-DNA insertion point in the line Salk-103699 (*glxI;2*), Primer binding sites and potential sgRNA target sites. black boxes = exons; grey boxes = untranslated regions; green boxes = sgRNA target sites. Triangle = T-DNA insertion point. **B,** PCR with primers pCAS@glxI;2_F and pCAS@glxI;2_R to verify the insertion of the pUB-Cas9-T-DNA Insertion in *A. thaliana* Col-0. The expected PCR fragment has a length of 1 kb. All nine plants resulted positive for the insertion and thus named CRISPR/Cas9@glxI;2-# (CC@glxI;2-#). **C,** PCR with primers F07+R09 to reveal potential deletions between both sgRNA1 and 2. Wild type plants present a 770 bp fragment. Plant line CC@glxI;2-1 showed a smaller amplification band, indicating the loss of a gene fragment. Sequencing of the PCR fragment confirmed the deletion. **D,** Digestion of PCR fragments obtained with primers F08+R08 with Bsu36I to reveal point mutations at sgRNA1. The wild-type fragment of 490 bp is cleaved into two fragments of 387 and 100 bp. Resistance to restriction in CC@glxI;2-1, -3, -6, and -8 indicate the presence of a point mutation. **E,** Digestion of PCR fragments obtained with primers F09+R09 with StyI to reveal point mutations at sgRNA2. The wild-type fragment of 448 bp is cleaved into two fragments of 266 and 182 bp. Resistance to restriction in CC@glxI;2-1, -2, -3, -5, -6, -8, and -9 indicates point mutations at the target site. All primers used are listed in Supplemental Table 1.

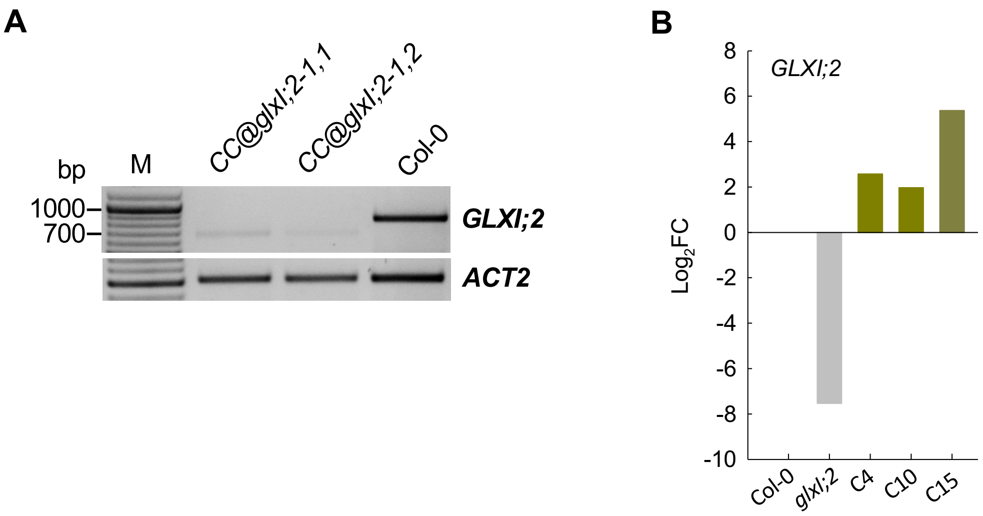


**Supplemental Figure 7. Analysis of *GLXI;2* expression in Arabidopsis *GLXI;2* loss of function and complemented lines. A,** Semi-quantitative PCR showing low expression of a shortened GLXI;2 transcript in two plants of the CC@glxI;2-1 line. The expected wild-type PCR fragment has 852 bp, that bearing the deletion has 686 bp. M= Molecular weight markers. *ACTIN2 (ACT2)* was used as reference. **B,** Relative change of *GLXI;2* transcript abundance in *glxI;2* and the complemented lines C4, C10, and C15 relative to the wild type (Col-0) measured through quantitative real time PCR. *ACTIN2* was used as reference.

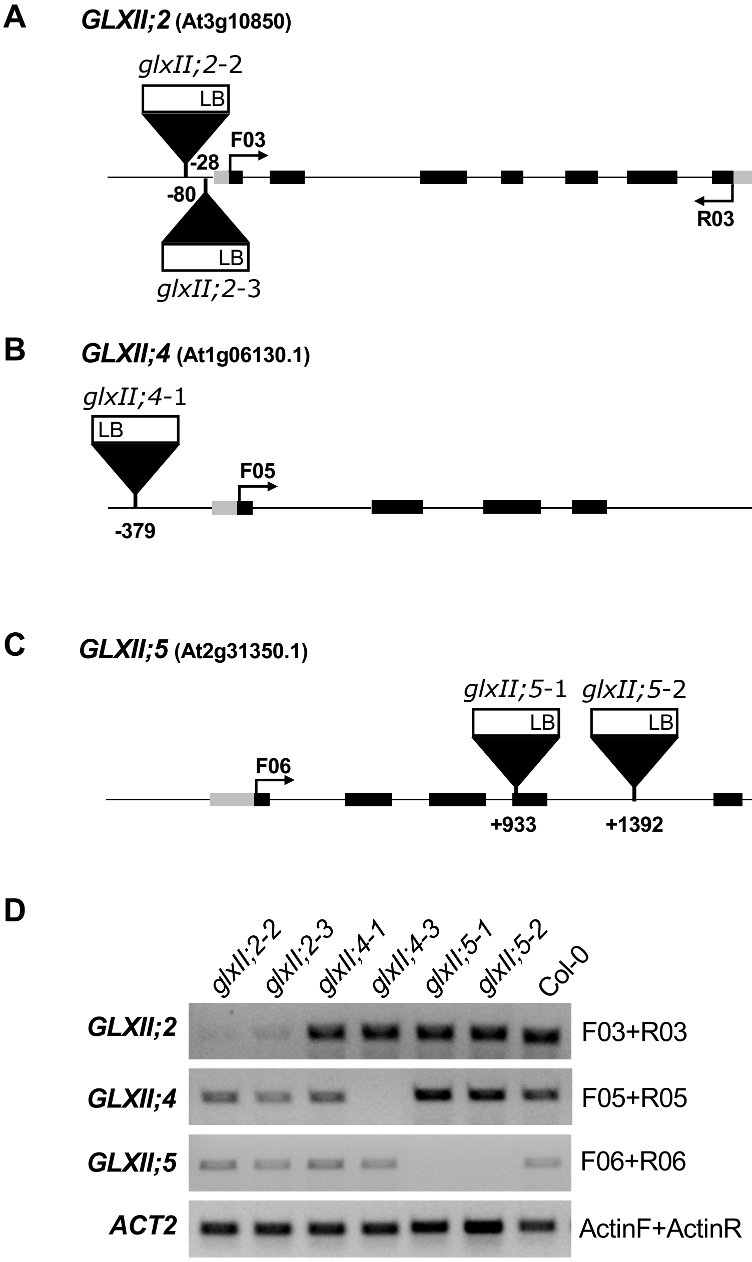


**Supplementary Figure 8. Arabidopsis GLXII T-DNA insertion lines. A,** Schematic representation of the *GLXII;2* gene locus showing the verified T-DNA insertion points in lines Salk_134008 (*glxII;2-2*) and Salk_090826 (*glxII;2-3*) and the primer pair F03+R03 used for the amplification of *GLXII;2* transcripts via RT-PCR. **B,** Schematic representation of the *GLXII;4* gene locus showing the verified T-DNA insertion points in lines Flag_603A03 (*glxII;4-1*), and Salk_045876 (*glxII;4-3*) and the primer pair F04+R04 used for the amplification of *GLXII;4* transcripts via RT-PCR. **C,** Schematic representation of the *GLXII;5* gene locus showing the verified T-DNA insertion points in lines Salk_014157 (*glxII;5-1*) and Salk_100618 (*glxII;5-2*) and the primer pair F05+R05 used for the amplification of *GLXII* transcripts via RT-PCR. **D,** Amplification of *GLXII* transcripts via RT-PCR using the primers shown on the right (Suppl. Table 1) *ACTIN2* was used as reference. box = exon; black = coding sequence; grey = untranslated region; Triangle = T-DNA insertion point.

**Supplementary Table 1.** Sequence of the primers used in this study. Primer sequence is 5' to 3'. * = primer taken from Czechowski et al., 2005, F = forward, R = reverse.

| Purpose | Name | Sequence 5'-3' |
|---|---|---|
| Expression of GLXI;2 | OX-GLXI;2_F | CATCGATAGTACTGTCGACCAACAGACGGTTTACAGAAGC |
| | OX-GLXI;2_R | TCTTCATCTTCATATGAGCTACAATCAAAATTGGTCCGCA |
| GLXI;2 cloned in pUBQ10 | pHyg_F | CGTGATCAAGGTAAATTTCTGTGT |
| | MS_gly1-2_R | TTGAGTTGAAATGGCAAAATGCCCA |
| T-DNA screening | Salk_LB | TGGTTCACGTAGTGGGCCATCG |
| | koGLXII;2-3_F | AGCCACCAATCAAACAATGAG |
| | koGLXII;2-3_R | TCAGAGACCTTGTGGTGGATC |
| | koGLXII;4-1_F | CCCATTACAATTTTGCAATGG |
| | koGLXII;4-1_R | TTGGAGGAAATGATTCAGCAC |
| | koGLXII;4-3_F | CATTTTGAGCCTCAAGAATTTG |
| | koGLXII;4-3_R | ACGTAACGTCACGGTCAAAAC |
| | koGLXII;5-1_F | TTGAGAAAGCTTCTTGTTGCC |
| | koGLXII;5-1_R | ACGTTTAGATGTGCATTTGGC |
| | koGLXII;5-2_F | TTGGAATTACTCTGCACACCC |
| | koGLXII;5-2_R | ATTCCTGGAATTGATATGGCC |
| | koGLXII;2-2_F | AGCCACCAATCAAACAATGAG |
| | koGLXII;2-2_R | TCAGAGACCTTGTGGTGGATC |
| RT-PCR for GLXII T-DNA insertion lines | F03 | AGATCTTCCACGTTCCTTGTCT |
| | R03 | CCCCTCCACTGATCCTTCTTG |
| | F05 | ATGCAAGCCATCTCCAAAGT |
| | R05 | GTGTTCTCAGTGCGGAGAAAC |
| | F06 | TCTCGAAAGCTTCTTCTGCCA |
| | R06 | GCTTAGAAATCATCCTTTGCTTTCC |
| | Actin_F | TAACTCTCCCGCTATGTATGTCGC |
| | Actin_R | GAAGCAAGAATGGAACCACCG |
| GLXI;2 qPCR | RL_GLXI;2_F | AGGACCCTGATGGCTACACT |
| | RL_GLXI;2_R | CATCCCGAGGGCCTTTTCAT |
| | RL_Act_F* | CTTGCACCAAGCAGCATGAA |
| | RL_Act_R* | CCGATCCAGACACTGTACTTCCTT |
| GLXI;2 sgRNA | pFH6_1_GLXI;2_F | ATTGGCATTAGAATACTTCTCCTC |
| | pFH6_1_GLXI;2_R | AAACGAGGAGAAGTATTCTAATGC |
| | pFH6_2_GLXI;2_F | ATTGGAGAACGTCCGTGCCAAGGG |
| | pFH6_2_GLXI;2_R | AAACCCCTTGGCACGGACGTTCTC |
| CC@glxI;2 screening | pCAS@glxI;2-1_F | CTGCCCAACGAGAAGGTGC |
| | pCAS@glxI;2-2_R | AGTCGGACAGGCGGTTGATG |
| | F07 | GAAACATGGCTGAGGCTTCTGA |
| | F08 | CAGGAGAGAGCAAGTTTATAGGGGAA |
| | R08 | TAACAGGAGAGGGCGAGTAGAG |
| | F09 | AGTATGCACTCTACTCGCCCTC |
| | R09 | TTTCATAGAATTTGATGGCACG |
| RT-PCR for CC@glxI;2 | Gly-h-At1g11840 FW | CACCATGGCTGAGGCTTCTGATTTGTTGGAA |
| | Gly-h-At1g11840 RV | TCATTCCAGTTCCTTGAGAAAATCTTTGTTGTC |
