## Supplemental Dataset7. Statistic analysis for "Viridiplantae-specific GLXI and GLXII isoforms co-evolved and detoxify glucosone *in planta*"

**Supplementary File 7: Summary of statistics**

**Figure 4 (B)**

| Two-way ANOVA | Ordinary |  |  |  |
| --- | --- | --- | --- | --- |
| Alpha | 0,05 |  |  |  |
| Source of Variation | % of total variation | P value | P value summary | Significant? |
| Interaction | 20,26 | <0,0001 | **** | Yes |
| Genotype | 20,38 | <0,0001 | **** | Yes |
| Treatment | 55,81 | <0,0001 | **** | Yes |

**Figure 5 (B)**

| Two-way ANOVA | Ordinary |  |  |  |
| --- | --- | --- | --- | --- |
| Alpha | 0,05 |  |  |  |
| Source of Variation | % of total variation | P value | P value summary | Significant? |
| Interaction | 19,00 | <0,0001 | **** | Yes |
| Treatment | 63,22 | <0,0001 | **** | Yes |
| Genotyp | 15,72 | <0,0001 | **** | Yes |

**Figure 5 (C)**

0.1mM KDG

| Dunnett's multiple comparisons test | Significant? | Summary | Adjusted P Value |
| --- | --- | --- | --- |
| Col-0 vs. *glxI,2* | No | ns | 0,9659 |
| Col-0 vs. *C4* | No | ns | >0,9999 |
| Col-0 vs. *glxI;3* | No | ns | 0,9769 |
| Col-0 vs. *glxI;I* | No | ns | 0,9997 |
| Col-0 vs. C15 | No | ns | >0,9999 |
| Col-0 vs. CC@glxI,2 | No | ns | 0,9961 |

0.25 mM KDG

| Dunnett's multiple comparisons test | Significant? | Summary | Adjusted P Value |
| --- | --- | --- | --- |
| Col-0 vs. *glxI,2* | No | ns | 0,1165 |
| Col-0 vs. *C4* | No | ns | 0,9417 |
| Col-0 vs. *glxI;3* | No | ns | 0,5576 |
| Col-0 vs. *glxI;I* | No | ns | 0,5247 |
| Col-0 vs. C15 | No | ns | 0,4744 |
| Col-0 vs. CC@glxI,2 | No | ns | 0,9985 |

0.5 mM KDG

| Dunnett's multiple comparisons test | Significant? | Summary | Adjusted P Value |
| --- | --- | --- | --- |
| Col-0 vs. *glxI,2* | Yes | * | 0,0109 |
| Col-0 vs. *C4* | No | ns | 0,9649 |
| Col-0 vs. *glxI;3* | No | ns | 0,0802 |
| Col-0 vs. *glxI;I* | No | ns | 0,1168 |
| Col-0 vs. C15 | No | ns | 0,3623 |
| Col-0 vs. CC@glxI,2 | Yes | * | 0,0378 |

1 mM KDG

| Dunnett's multiple comparisons test | Significant? | Summary | Adjusted P Value |
| --- | --- | --- | --- |
| Col-0 vs. *glxI,2* | Yes | *** | 0,0002 |
| Col-0 vs. *C4* | No | ns | 0,9745 |
| Col-0 vs. *glxI;3* | No | ns | 0,0514 |
| Col-0 vs. *glxI;I* | Yes | * | 0,0284 |
| Col-0 vs. C15 | No | ns | 0,0864 |
| Col-0 vs. CC@glxI,2 | Yes | ** | 0,0053 |

2 mM KDG

| Dunnett's multiple comparisons test | Significant? | Summary | Adjusted P Value |
| --- | --- | --- | --- |
| Col-0 vs. *glxI,2* | Yes | *** | 0,0006 |
| Col-0 vs. *C4* | No | ns | 0,9732 |
| Col-0 vs. *glxI;3* | No | ns | 0,0891 |
| Col-0 vs. *glxI;I* | No | ns | 0,0850 |
| Col-0 vs. C15 | No | ns | 0,1927 |
| Col-0 vs. CC@glxI,2 | Yes | *** | 0,0004 |

**Figure 6 (B)**

| Two-way ANOVA | Ordinary |  |  |  |
| --- | --- | --- | --- | --- |
| Alpha | 0,05 |  |  |  |
| Source of Variation | % of total variation | P value | P value summary | Significant? |
| Interaction | 3,340 | <0,0001 | **** | Yes |
| Treatment | 86,61 | <0,0001 | **** | Yes |
| Genotype | 4,924 | <0,0001 | **** | Yes |

**Figure 7 (D)**

| Dunnett's multiple comparisons test | Significant? | Summary | Adjusted P Value |
| --- | --- | --- | --- |
| Col-0 vs. *glxI;2* | Yes | **** | <0,0001 |
| Col-0 vs. *CC@glxI;2* | Yes | **** | <0,0001 |
| Col-0 vs. *C4* | No | ns | 0,9997 |
| Col-0 vs. *C15* | No | ns | 0,9171 |
| Col-0 vs. *glxII;4-3/5-2* | No | ns | 0,9989 |

**Figure 7 (E)**

| Dunnett's multiple comparisons test | Significant? | Summary | Adjusted P Value |
| --- | --- | --- | --- |
| Col-0 vs. glxI;2 | Yes | **** | <0,0001 |
| Col-0 vs. CCglxI;2 | Yes | **** | <0,0001 |
| Col-0 vs. C4 | No | ns | 0,3555 |
| Col-0 vs. C15 | No | ns | 0,9688 |
| Col-0 vs. glxII,4-3/5-2 | Yes | *** | 0,0001 |

**Figure 8 (B)**

| Column B | Col-0 2%Glc |
| --- | --- |
| vs. | vs, |
| Column A | Col-0 MS |
| Unpaired t test |  |
| P value | 0,0002 |
| P value summary | *** |
| Significantly different (P < 0.05)? | Yes |

**Figure 8 (C)**

| Column B | Col-0 2%Glc |
| --- | --- |
| vs. | vs, |
| Column A | Col-0 MS |
| Unpaired t test |  |
| P value | 0,0095 |
| P value summary | ** |
| Significantly different (P < 0.05)? | Yes |
